# Powdery mildew effectors AVR_A1_ and BEC1016 target the ER J-domain protein *Hv*ERdj3B required for immunity in barley

**DOI:** 10.1101/2022.04.27.489729

**Authors:** Zizhang Li, Valeria Velásquez-Zapata, J. Mitch Elmore, Xuan Li, Wenjun Xie, Sohini Deb, Xiao Tian, Sagnik Banerjee, Hans J. L. Jørgensen, Carsten Pedersen, Roger P. Wise, Hans Thordal-Christensen

**Affiliations:** Department of Plant and Environmental Sciences, University of Copenhagen, Thorvaldsensvej 40, 1871 Frederiksberg C, Denmark; Program in Bioinformatics & Computational Biology, Iowa State University, Ames, IA, 50011, USA; Department of Plant Pathology, Entomology and Microbiology, Iowa State University, Ames, IA, 50011 USA; Department of Statistics, Iowa State University, Ames, IA, 50011, USA; USDA-Agricultural Research Service, Corn Insects and Crop Genetics Research Unit, Ames, IA, 50011, USA

**Keywords:** Barley, powdery mildew, immunity, pathogen effectors, Yeast two-hybrid Next-generation interaction screening, ER quality control, ERdj3b, signal peptide-independent ER-uptake

## Abstract

The barley powdery mildew fungus, *Blumeria hordei* (*Bh*), secretes hundreds of candidate secreted effector proteins (CSEPs) to facilitate pathogen infection and colonization. One of these, CSEP0008, is directly recognized by the barley nucleotide-binding leucine-rich-repeat (NLR) receptor, MLA1, and therefore designated AVR_A1_. Here we show that AVR_A1_ and the sequence-unrelated *Bh* effector BEC1016 (CSEP0491) suppress immunity in barley. We used yeast two-hybrid next-generation interaction screens (Y2H-NGIS), followed by binary Y2H and *in planta* protein-protein interactions studies, and identified a common barley target of AVR_A1_ and BEC1016, the endoplasmic reticulum (ER)-localized J-domain protein, *Hv*ERdj3B. Silencing of this ER quality control (ERQC) protein increased the *Bh* penetration. *Hv*ERdj3B is ER luminal, and we showed using split GFP that AVR_A1_ and BEC1016 translocate into the ER - signal peptide-independently. Silencing of *Hv*ERdj3B and expression the two effectors hampered trafficking of a vacuolar marker through the ER as a shared cellular phenotype, agreeing with the effectors targeting this ERQC component. Together, these results suggest that the barley innate immunity, preventing *Bh* entry into epidermal cells, is dependent on ERQC, which in turn requires the J-domain protein, *Hv*ERdj3B, regulated by AVR_A1_ and BEC1016. Plant disease resistance often occurs upon direct or indirect recognition of pathogen effectors by host NLR receptors. Previous work has shown that AVR_A1_ is directly recognized in the cytosol by the immune receptor, MLA1. We speculate that the AVR_A1_ J-domain target being inside the ER, where it is inapproachable by NLRs, has forced the plant to evolve this challenging direct recognition.

**SIGNIFICANCE:** The complex plant immune system is highly dependent on fundamental cellular machineries, such as the endomembrane system and the ER quality control (ERQC), essential for delivery of immunity-associated membrane-bound and endomembrane soluble proteins to their destinations. We now find that pathogen effectors can interact with an ERQC component and suppress immunity, thereby adding to the molecular insight in plant-pathogen interactions.

## INTRODUCTION

Plants have evolved a multilayered and interconnected innate immune system to protect themselves against pathogen attacks (Ngou et al., 2022; Yuan et al., 2021a). An initial layer, designated pathogen-associated molecular <u>p</u>attern-<u>t</u>riggered immunity (PTI) is activated by <u>p</u>attern-<u>r</u>ecognition <u>r</u>eceptors (PRRs) located on the <u>p</u>lasma <u>m</u>embrane (PM) (Zipfel & Oldroyd, 2017). However, plant pathogens can secrete large numbers of effectors into the host to interfere with many host cellular processes in order to suppress PTI and establish parasitism (Figueroa et al., 2021; Kanja & Hammond-Kosack, 2020;). In response, plants have evolved <u>e</u>ffector-triggered immunity (ETI) that is initiated upon direct or indirect recognition of effectors by specific intracellular <u>n</u>ucleotide-binding leucine-rich <u>r</u>epeat (NLR) receptors (Jones et al., 2016; Ngou et al., 2022; Thordal-Christensen, 2020). With some exceptions, NLR receptors represent the most utilized class of resistance proteins in agriculture (Kourelis & van der Hoorn, 2018; Ngou et al., 2022; Sun et al., 2020; van Wersch et al., 2020). These innate immune systems rely on basic cellular processes, such as protein quality control systems. An intact <u>e</u>ndoplasmic <u>r</u>eticulum (ER) <u>q</u>uality <u>c</u>ontrol (ERQC) is required to transport certain *de novo* synthesized PRRs through the ER for them to be trafficked to their destination at the PM (Tintor & Saijo, 2014). For example, trafficking of the Arabidopsis <u>EF</u>-Tu <u>r</u>eceptor, EFR, is particularly dependent on the ERQC, where it requires the chaperones, <u>ER D</u>naJ3B protein (ERdj3B) and <u>b</u>inding immunoglobulin <u>p</u>rotein (BiP), as well as <u>s</u>tromal cell-<u>d</u>erived factor 2 (SDF2), for its proper folding and glycosylation (Nekrasov et al., 2009). While ERdj3B is an ER luminal J-domain protein, homologous proteins elsewhere in the cell have been implicated in plant immunity as well. For instance, in rice infected by the blast fungus, *Magnaporthe oryzae*, cytosolic and mitochondria-associated J-domain proteins are essential for immunity, and interestingly, the latter protein is targeted by the disease-promoting fungal effector, *Mo*CDIP4 (Xu et al., 2020a; Zhong et al., 2018). J-domain proteins are related to the bacterial DnaJ chaperones, also known as heat shock proteins 40 (HSP40). They interact with HSP70 proteins, such as BiP in the ER, to which they deliver misfolded client proteins and stimulate the HSP70 ATPase activity (Fatima et al., 2021; Pobre et al., 2019).

Pathogen effectors manipulate diverse aspects of host biology during infection (Speth *et al*. 2007). Several recent genomics, transcriptomics and proteomics studies have identified some 600 <u>c</u>andidates for <u>s</u>ecreted <u>e</u>ffector <u>p</u>roteins (CSEPs) in the barley powdery mildew fungus, *<u>B</u>lumeria <u>h</u>ordei* (*Bh*) (Bindschedler et al., 2009; Frantzeskakis et al., 2018; Menardo et al., 2017; Pedersen et al., 2012; Spanu et al., 2010). However, only a subset of these have been studied in detail. <u>H</u>ost-induced <u>g</u>ene <u>s</u>ilencing (HIGS) analyses have demonstrated that approximately 20 CSEPs contribute to *Bh* virulence, which is ∼25% of those studied (Aguilar et al., 2016; Ahmed et al., 2015, 2016; Li et al., 2021; Pliego et al., 2013; Yuan et al., 2021b; Zhang et al., 2012). Moreover, Li et al. (2021) screened ∼100 CSEPs from *Bh* and found 15 of them to suppress BAX-induced programmed cell death in *Nicotiana benthamiana*. Two of those, CSEP0139 and CSEP0182, also suppressed BAX-induced programmed cell death in barley. To date, only a few plant targets of *Bh* effectors have been identified. CSEP0055 targets the barley defense proteins, PR1 and PR17 (Zhang et al., 2012), while CSEP0105 and CSEP0162 target the barley small heat shock proteins, Hsp16.9 and Hsp17.5 (Ahmed et al., 2015). CSEP0064 and CSEP0264 target PR10, while CSEP0264, historically designated <u>B</u>lumeria <u>e</u>ffector <u>c</u>andidate (BEC)1054, also targets eukaryotic elongation factor1 alpha (Pennington et al., 2019). Likewise, CSEP0027 was recently found to target a barley catalase (Yuan et al., 2021b).

In addition to virulence functions, six *Bh* CSEPs are directly recognized as avirulence proteins by NLRs encoded by alleles of the *Mla* powdery mildew resistance locus (Bauer et al., 2021; Lu et al., 2016; Saur et al., 2019, Seeholzer et al., 2010). For instance, CSEP0008 (gene ID BLGH_03023) is recognized by MLA1, and thus named AVR_A1_ (Lu et al., 2016). Other recognized CSEPs are CSEP0059 (AVR_A7_), CSEP0174 (AVR_A9_), CSEP0141 (AVR_A10_ and AVR_A22_), and CSEP0372 (AVR_A13_) (Bauer et al., 2021; Cao et al., 2023; Lu et al., 2016; Saur et al., 2019;). AVR_A6_ is represented by three near-identical copies in the DH14 genome, BLGH_00709 (CSEP0254), BLGH_00708 and BLGH_07091 (Bauer et al., 2021; Cao et al., 2023), which may be expressed in an isolate-specific manor (Velásquez-Zapata et al., 2023b).

Early studies have shown that effectors screened against immune-associated proteins of Arabidopsis identified several ‘hub’ proteins that are targeted by multiple effectors (Mukhtar et al. 2011; Weßling et al. 2014). To test this observation in the barley-*Bh* system, we initiated a screen for protein-protein interactions with several CSEPs, focusing on cloned AVR effectors (Lu et al., 2016), as well as those that displayed unique transcript abundance during infection and/or a significant HIGS phenotype (Pliego et al. 2013). Here we benefit from recent advances in <u>y</u>east two-<u>h</u>ybrid (Y2H) analyses, termed <u>n</u>ext-<u>g</u>eneration interaction <u>s</u>creening (NGIS) (Suter et al. 2015), which uses deep sequencing to score the output from Y2H screens (Erffelinck et al., 2018; Lewis et al., 2012, Pashkova et al., 2016; Trigg et al., 2017; Weimann et al., 2013; Yachie et al., 2016). These approaches facilitate a quantitative measure of which preys interact with each bait protein (Suter et al. 2015) and identify reproducible <u>p</u>rotein-<u>p</u>rotein interactions (PPI) with 70-90% accuracy (Pashkova et al., 2016; Trigg et al., 2017; Velásquez-Zapata et al., 2021; Weimann et al., 2013). This then allows functional PPI to be positioned within interactome networks (Mukhtar et al., 2011; Velásquez-Zapata et al., 2021, 2022; Weßling et al., 2014).

In our NGIS pipeline, we found that AVR_A1_ (CSEP0008, gene ID BLGH_03023) and BEC1016 (CSEP0491, gene ID BLGH_07006), which has not been shown to be an avirulence protein, both target the barley J-containing protein, *Hv*ERdj3B. As indicated, AVR_A1_ is recognized as an avirulence protein by barley MLA1, but no virulence contribution was found for it (Lu et al., 2016). In contrast, BEC1016 has no documented avirulence function, but it contributes significantly to virulence (Pliego et al. 2013). Using a split GFP system, our analyses confirmed that *Hv*ERdj3B is an ER-luminal protein and we could show that both effectors translocate <u>s</u>ignal <u>p</u>eptide (SP)-independently from the plant cytosol into the ER lumen. Silencing *Hv*ERdj3B, as well as overexpression of AVR_A1_ and BEC1016, not only enhanced the formation of fungal haustoria as an immunity-related phenotype, it also hampered trafficking of a vacuolar marker protein through the ER as a shared cellular phenotype. Together, our results indicate an essential role of ERQC in barley innate immunity and suggest that *Bh* effectors can interfere with ERQC processes.

## RESULTS

### AVR_A1_ and BEC1016 suppress immunity

The two effector candidates AVR_A1_ (singleton) and BEC1016 (CSEP family 5) (Pedersen et al., 2012) target the same barley protein (see below). They belong to the superfamily of <u>R</u>N<u>a</u>se-like <u>p</u>roteins associated with <u>h</u>austoria (RALPH) effectors with features resembling catalytically inactive RNases (Spanu, 2017), but their amino acid sequences are very different (Figure S1). They exhibit similar patterns of transcript accumulation in incompatible and compatible interactions up to *Bh* penetration of epidermal cells. However, they diverge during development of haustoria, when in particular the *AVR_A1_* transcript has a notable increase in the compatible interaction (Figure S2a,b). As an avirulence protein recognized by MLA1, AVR_A1_ is predicted also to have virulence function. However, silencing of *AVR_a1_* did not result in a significant decrease in haustoria in the original HIGS assay, unlike the case for BEC1016 (Pliego et al., 2013). As an alternative approach to study virulence functions, we transiently over-expressed AVR_A1_ and BEC1016 (w/o <u>s</u>ignal <u>p</u>eptide, SP) in leaf epidermal cells of one-week-old Golden Promise (susceptible) barley plants using particle bombardment. Two days later, the leaves were inoculated with *Bh* isolate C15, and after another 2 days transformed cells were scored for presence of haustoria, as evidence for successful penetration and fungal infection. Our data revealed that over-expression of AVR_A1_ and BEC1016 resulted in 96% and 150% higher haustoria numbers, respectively. However, only the latter effect was statistically significant and reflected an increased penetration rate (Figure 1a).

**Figure 1.**
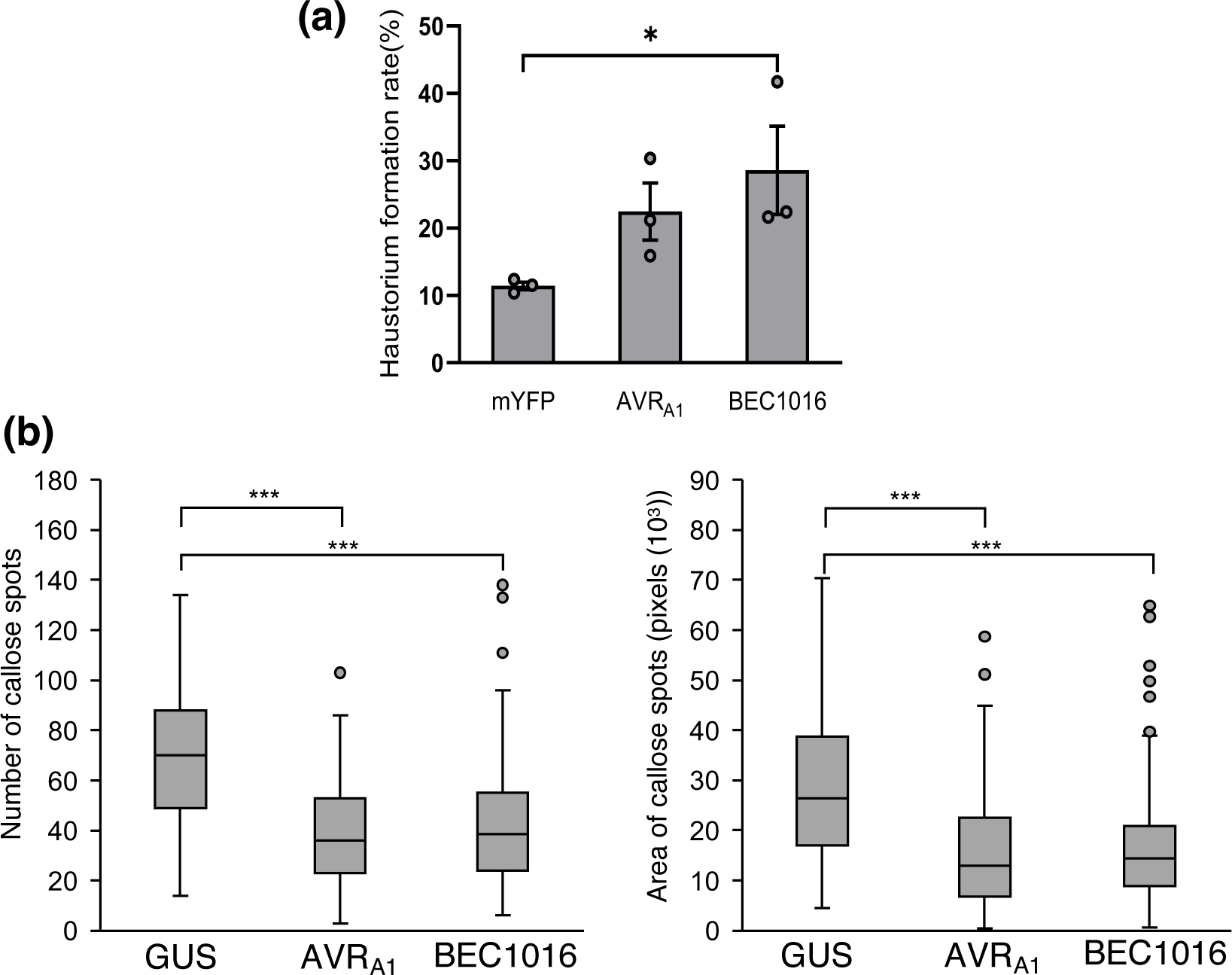
Influence of AVR_A1_ and BEC1016 on *Bh* infection and callose deposition in barley. (a) Effect of transient over-expression of AVR_A1_ and BEC1016 on *Bh* invasion. pUbi promoter-driven over-expression constructs were co-bombarded with a GUS-reporter constructs into leaf epidermal cells of 1-week-old barley and 2 days later inoculated with *Bh*. After another 2 days, the fungal haustorium formation was scored in GUS-expressing cells. Data shown are mean values of three independent experiments. Error bars, SE. *, *P* = 0.0179 calculated using a logistic regression model. (b) AVR_A1_ and BEC1016 can reduce bacterium-triggered callose deposition in barley plants estimated as numbers of callose spots (left) and area of callose spots (right). Leaves of eight-day-old barley plants were infiltrated with EtHAn, transformed with pEDV6-GUS, pEDV6-AVR_A1_ or pEDV6-BEC1016, and 24 h later the callose deposition we scored. Boxplots for the total number and total area of callose depositions show data for all pictures taken across biological replicates under each treatment. A repeated measures ANOVA followed for a paired sample t test were used to compare the treatments. ***, *P* < 2×10^-16^. Data shown are based on five independent experiments, statistically treated as replicates.

Pathogen-induced callose depositions function as a chemical and physical reinforcement of the plant cell wall towards invading pathogens (Voigt, 2014). To determine whether AVR_A1_ and BEC1016 affect callose deposition, they were introduced into barley leaf cells using EtHAn, a strain of *Pseudomonas fluorescens* modified to express the type-3 secretion system (T3SS) (Thomas et al., 2009; Upadhyaya et al., 2014). To verify protein transfer to the barley cells, EtHAn transformed with a construct expressing β-<u>glu</u>coronida<u>s</u>e (GUS) fused to a signal peptide for T3SS resulted in clear and uniform blue staining inside the leaf cells after reaction buffer incubation. There was no sign of stained bacteria in the apoplast (Figure S3), indicating GUS to be efficiently transferred into the barley cell. While EtHAn itself triggers callose formation, it can at the same time be studied how this is influenced by effectors (Sohn et al., 2007; Xu et al., 2020b). Five replicates of eight-day-old Golden Promise first leaves were infiltrated with EtHAn expressing GUS (negative control), AVR_A1_ or BEC1016. After 24 h, both the total number and total area of callose depositions were significantly reduced by the effectors (Figure 1b). The number of bacteria after 24 h was the same for the three strains (Figure S4), supporting that the effectors directly influence the callose deposition. Taken together, it appears that AVR_A1_ and BEC1016 target and inhibit PTI, and thereby promote *Bh* penetration.

### Both AVR_A1_ and BEC1016 interact with barley J-domain protein, *Hv*ERdj3B

To identify host-pathogen PPI, three independent replications of batch Y2H-NGIS were performed according to Elmore et al. (2023) and Velásquez-Zapata et al. (2021, 2023a). The screens involved parallel histidine selection (for enrichment of yeast cells with bait and prey interactions) vs. non-selected controls (selection for bait and prey, but not their interactions). Deep Illumina sequencing was performed across the GAL4 activation domain (AD) and into the prey protein coding sequences for all cultures. Two complementary software packages, designated NGPINT and Y2H-SCORES, were used to quantify the outcome of Y2H-NGIS (Banerjee et al., 2021; Velásquez-Zapata et al., 2021, 2023a). NGPINT facilitates the identification of GAL4 AD-prey *in-frame* coding sequence fusion reads. Then Y2H-SCORES uses appropriate normalization methods, count data and statistical models to build a set of ranking scores to predict three properties expected from true interactors, *i.e.*, enrichment in selection vs. non-selection conditions, specificity to a bait screening and *in-frame* selection of the prey (Velásquez-Zapata et al., 2021). Prey proteins that interact with the selected baits, as indicated by significant enrichment under selection, are reconstituted *in silico* with their mapped fusion reads to identify the prey sequence containing the interaction domain for further studies. Note that this approach delineates the specific interaction region, and not necessarily the full-length clone, which is inferred from the reference genome annotation.

Using the AVR_A1_ and BEC1016 effectors as baits to screen a 3-frame Y2H library made from our infection time course (see Experimental Procedures), we identified HORVU1Hr1G022990, herein designated *Hv*ERdj3B (GenBank ID: AK376215.1 and KAE8793706.1) as a candidate interactor for both (Figure 2, S5). Ranking (Table S1) and visualization of the candidate interactors for each bait identified a prey fragment covering the C-terminal two-thirds of *Hv*ERdj3B (aa 142-350). Subsequently, binary Y2H according to Dreze et al. (2010) confirmed interaction between this *Hv*ERdj3B fragment with AVR_A1_ and BEC1016 (Figure S6).

**Figure 2.**
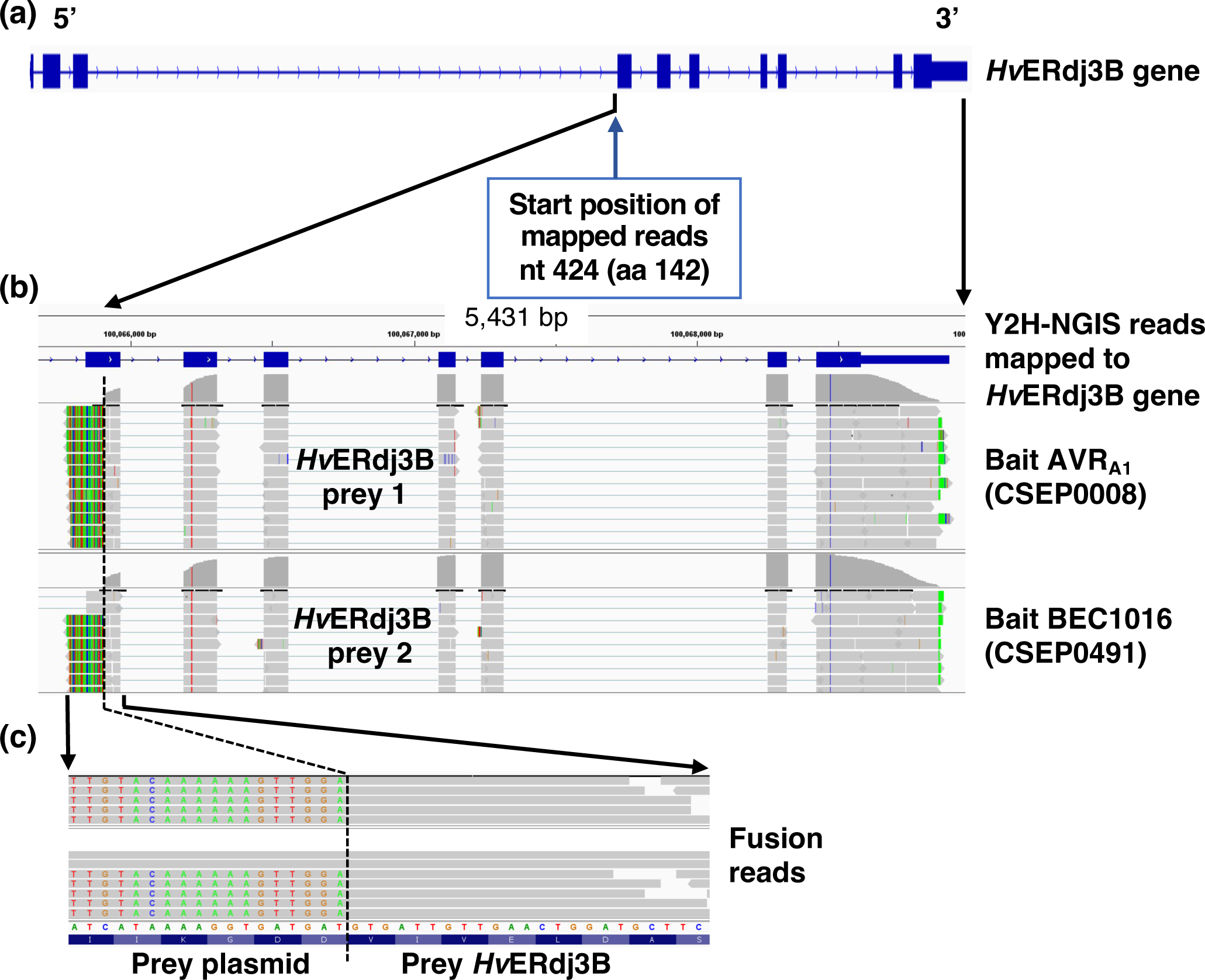
Y2H-NGIS used to identify interactions between AVR_A1_, BEC1016 and *Hv*ERdj3B. (a) *Hv*ERdj3B gene model with exons depicted as blue boxes and introns as hash marks. (b) Integrative genomic view (IGV) obtained from the software NGPINT after Y2H-NGIS analysis of the *Hv*ERdj3B prey reads selected by both the AVR_A1_ and the BEC1016 bait. Prey fragments were reconstructed from the reads mapped to the *HvERdj3B* gene (shown in grey across the exons) in each Y2H-NGIS dataset and located towards the 3’ end of the gene in both cases. (c) Expanded view of the 5’ fusion reads that allowed the determination of the frame and nucleotide-resolved prey fragments. The fusion reads contain a prey plasmid sequence (shown in different colors as mismatches from the reference gene) and the prey sequence. See also Fig. S5.

To study the specificity of these interactions further, we performed extensive binary Y2H assays with an independent colorimetric Y2H system (see Methods). The two effectors (w/o SP) were fused to either the Gal4 transcription factor DNA-<u>b</u>inding <u>d</u>omain (BD) or transcription-<u>a</u>ctivation <u>d</u>omain (AD) and combined with either the full-length (w/o SP), the C-terminal two-thirds (aa 142-350) or the central part (aa 142-231) of *Hv*ERdj3B fused to either the AD or the BD. In both orientations of the setup, both effectors were found to interact with the C-terminal two-thirds and the central part, but not the full-length of *Hv*ERdj3B (Figure 3). It is common to recover only a fragment and not a full-length protein interaction in Y2H assays. Several factors like protein size, domain toxicity and accessibility can explain a negative interaction in yeast, which may vary when tested *in planta* (Galletta & Rusan, 2015). Nevertheless, these results suggest that the N-terminus of *Hv*ERdj3B is not necessary for effector binding and that the interaction occurs only at the central part of *Hv*ERdj3B. In turn, this explains why reads did not appear from the 5’ end of the coding sequence in the Y2H-NGIS sequencing results (Figure 2).

**Figure 3.**
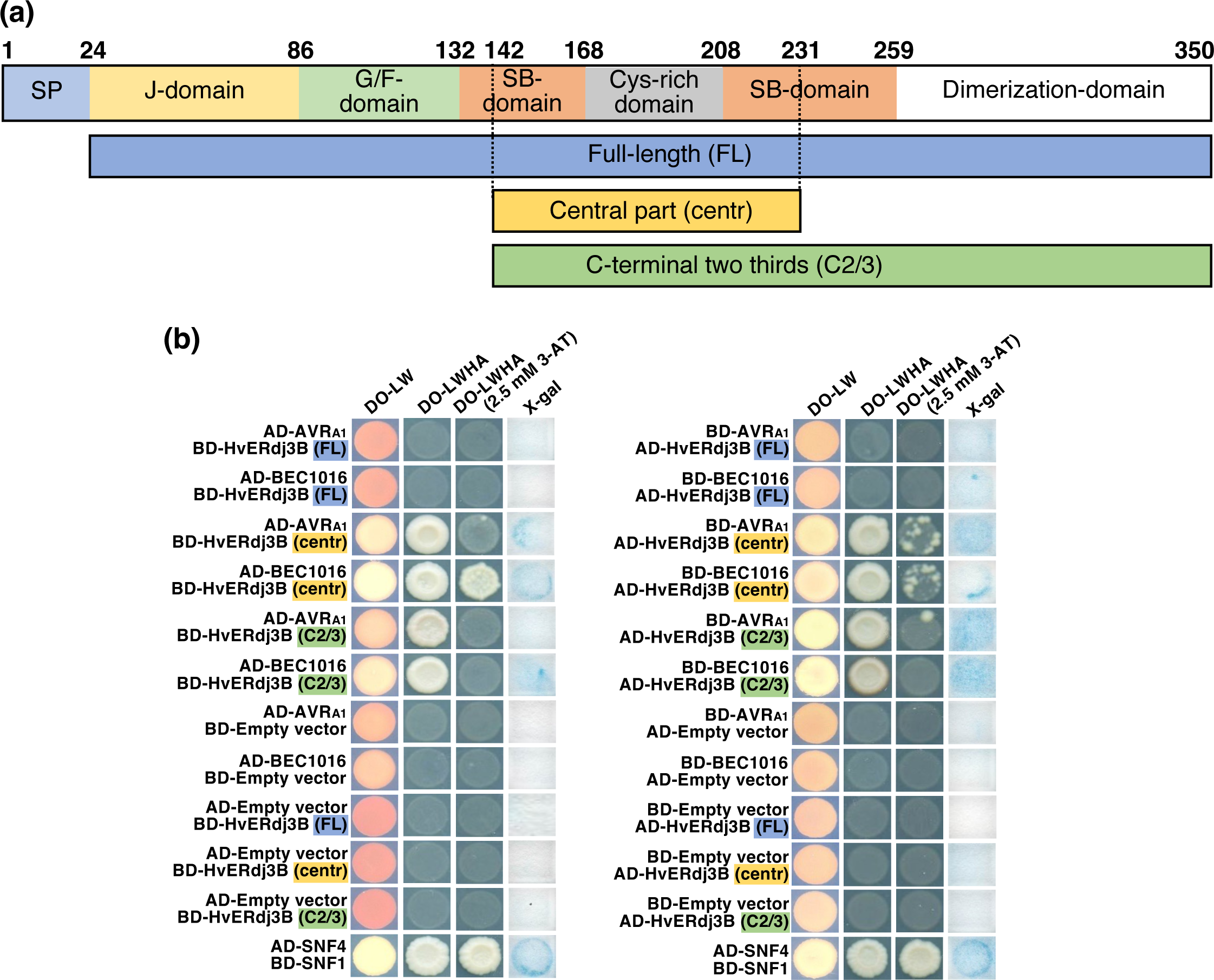
AVR_A1_ and BEC1016 interact with the central part of barley *Hv*ERdj3B in Y2H. (a) Domain structure of the 350 aa *Hv*ERdj3B protein and the Y2H-fragments. The domains are predicted according Chen, et al. (2017)(see **Fig. S5**). SP, signal peptide; G/F, glycine/phenylalanine; SB, substrate binding. Below the three Y2H-fragments are shown. (b) Yeast transformed with the destination vectors pDEST-AS2-1 [GAL4 binding-domain (BD)] and pDEST-ACT2-1 [GAL4 activation-domain (AD)] (Robertson, 2004) in different combinations of domain-fusions to effectors and the full-length (FL), the central part (centr) and the C-terminal two-thirds (C2/3) of *Hv*ERdj3B. Growth on <u>d</u>rop<u>o</u>ut (DO)-medium lacking leucine (L) and tryptophan (W) indicated presence of both constructs. Growth on DO-medium lacking L, W, histidine (H) and adenine (A) +/-2.5 mM <u>3</u>-<u>a</u>mino-1,2,4-triazole (3AT) indicated protein-protein interaction. β-galactosidase assay (X-gal), also indicating protein-protein interaction, was performed on filter paper prints of DO-LW-grown yeast. SNF1/SNF4 was used as a positive control and empty vectors were used as negative control.

To confirm interactions between AVR_A1_ or BEC1016 and *Hv*ERdj3B *in planta*, a <u>bi</u>molecular fluorescence <u>c</u>omplementation (BiFC) assay (Hu & Kerppola, 2003) was carried out in *N. benthamiana* leaf epidermal cells after Agrobacterium-mediated transformation. In this study, only the full-length *Hv*ERdj3B (w/o signal peptide) was used. Fluorescence was observed only when the *Hv*ERdj3B-cGFP fusion was combined with nGFP fused to the N-terminal or the C-terminal of the effectors (Figure 4a, S7). The specificity of the interactions was further documented by co-immunoprecipitation (co-IP) (Figure 4b).

**Figure 4.**
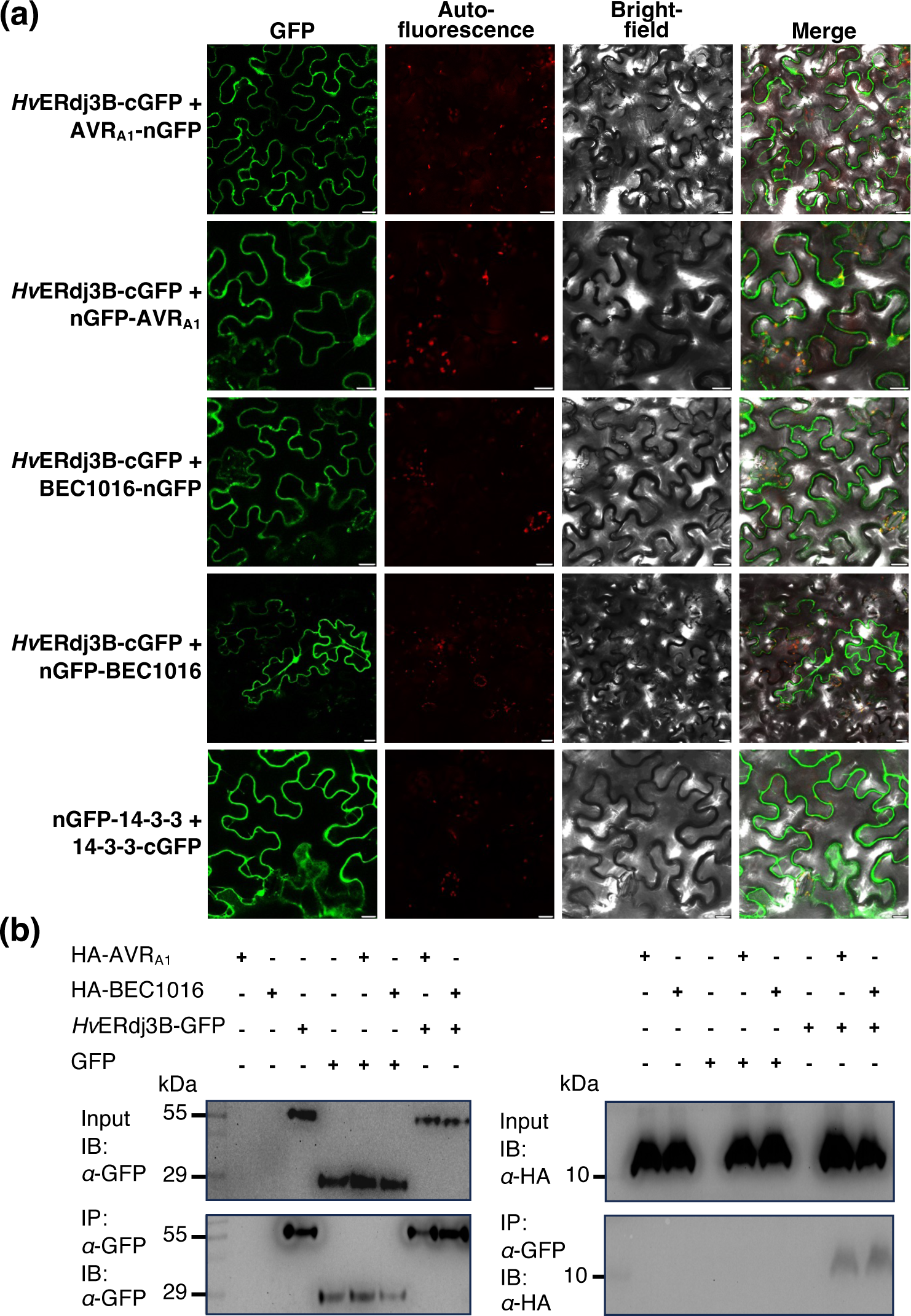
Interactions of *Hv*ERdj3B and AVR_A1_ as well as BEC1016 confirmed in living plant cells. (a) Reconstitution of fluorescent GFP (bi-fluorescence complementation) was observed in *N. benthamiana* epidermal cells expressing the shown combinations of the C and N-terminal parts of GFP fused with *Hv*ERdj3B (w/o SP) and AVR_A1_/BEC1016 (w/o SP). The remaining combinations did not allow fluorescent GFP to reconstitute (see **Fig. S7**). 14-3-3 dimerization was used as positive control. Expression constructs were introduced using *Agrobacterium* infiltration, and epidermal cells were observed by laser scanning confocal microscopy 48 h later. Size bars, 20 μm. (b) Co-immunoprecipitation of *Hv*ERdj3B and AVR_A1_/BEC1016. Construct combinations for expression of the indicated proteins (all w/o SP) were introduced in leaves of *N. benthamiana* using *Agrobacterium* infiltration. Three days later the proteins were extracted from the leaves and analyzed by SDS-PAGE/immunoblotting (IB) with anti-GFP and anti-HA antibodies before and after immunoprecipitation (IP) using anti-GFP magnetic beads. Expected protein molecular weight: GFP, 27 kDa, HvERdj3B-GFP, 65 kDa, HA-AVRA1, 13 kDa and HA-BEC1016, 13 kDa.

Taken together, these results confirmed that aa 142-231 of the J-domain protein *Hv*ERdj3B interacts with the diverged RALPH effectors, AVR_A1_ and BEC1016. According to alignment with the most closely related mammalian J-domain protein, this section corresponds with the predicted substrate-binding and the Cys-rich domains (Chen et al., 2017; Figure S5). Moreover, this domain appears accessible for binding the full-length *Hv*ERdj3B *in planta*, which is a prerequisite for the interactions to be functionally relevant.

### *Hv*ERdj3B is an ER luminal J-domain protein required for immunity during powdery mildew attack

*HvERdj3B* encodes a J-domain protein with an N-terminal signal peptide for ER targeting. The mature protein shares 77% amino acid identity with its closest Arabidopsis homologue, *At*ERdj3B (Figure S5), which is localized to the ER (Yamamoto et al., 2008). Significant accumulation of the *HvERdj3B* transcript was observed during *Bh* penetration, peaking at 20 h after inoculation, which would be consistent with a role for it in immunity (Figure S2c). Indeed, *At*ERdj3B has previously been implicated in immunity in Arabidopsis, where it is required for triggering of responses to the bacterial elongation factor-Tu (Nekrasov et al., 2009). To investigate whether *Hv*ERdj3B contributes to the immunity to *Bh*, we silenced the *HvERdj3B* gene in barley plants using <u>t</u>ransient-induced <u>g</u>ene <u>s</u>ilencing (TIGS) (Douchkov et al. 2005). Two RNAi constructs were generated with different *HvERdj3B* fragments and, together with a GUS reporter construct, bombarded into single epidermal cells of leaves of one-week-old Golden Promise plants. Two days later, the leaves were inoculated with *Bh* isolate C15 and formation of fungal haustoria was assessed after another 2 days. Our data revealed that silencing of *HvERdj3B* significantly enhanced the penetration rate, as *HvERdj3B*-RNAi-1 and *HvERdj3B*-RNAi-2 in average tripled the number of haustoria (*P* < 0.001). The *Mlo* RNAi positive control reduced the number of haustoria by ∼40% (Figure 5a). This increase in the level of infection suggests that *Hv*ERdj3B plays an essential role in barley immunity.

**Figure 5.**
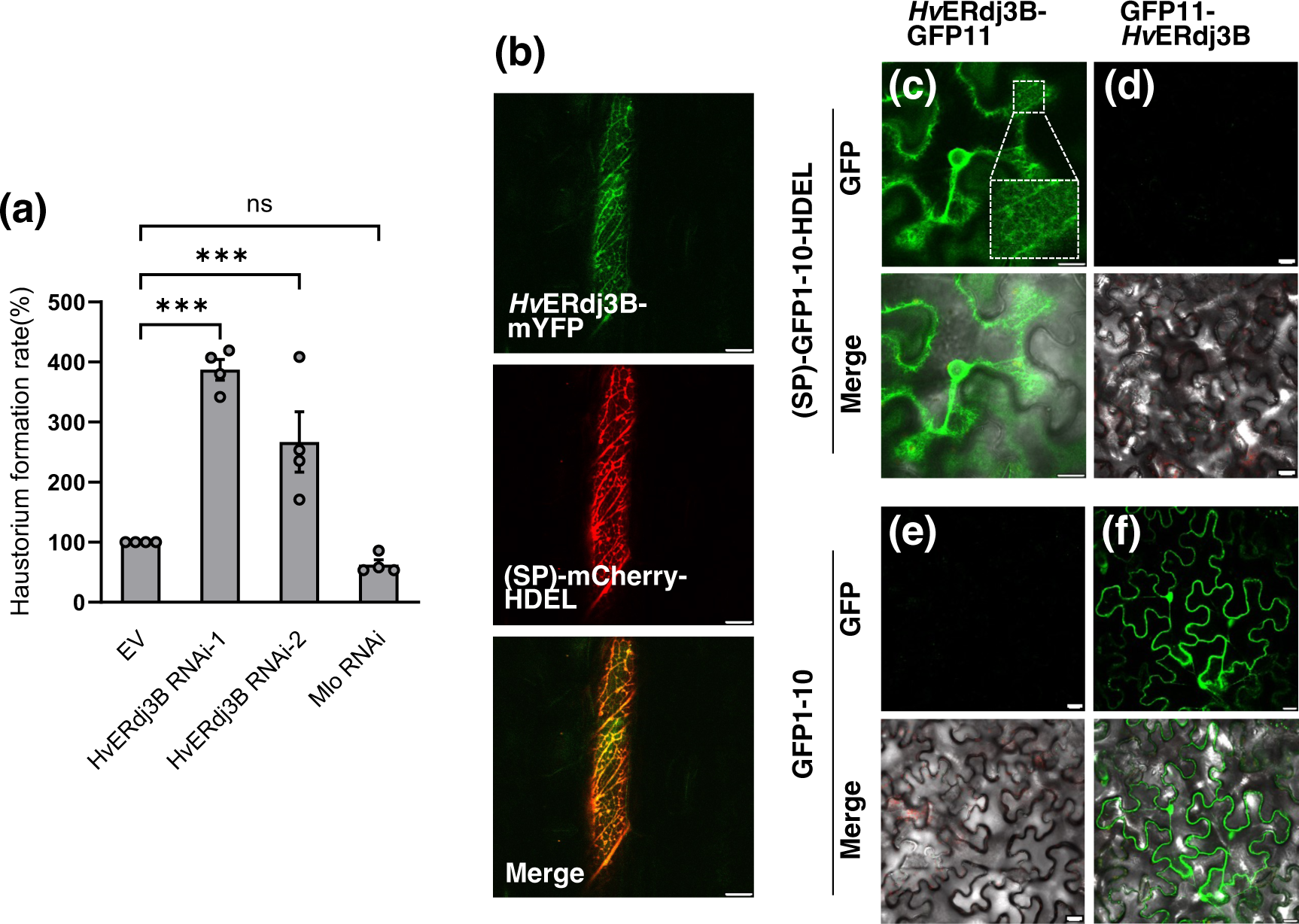
*Hv*ERdj3B is an ER luminal protein contributing to preinvasive immunity. (a) One-week-old barley leaves, bombarded with RNAi and GUS reporter constructs, were inoculated with *Bgh* and scored for fungal haustorium formation, calculated as the ratio of haustoria-containing transformed cells (GUS expressing cells) divided by the total number of transformed cells. Data shown are mean values of four independent experiments. Error bars, SE. ***, *P* < 0.001 calculated using a logistic regression model. (b) Localization of *Hv*ERdj3B in barley leaf epidermal cells. A ubiquitin promoter-driven expression construct encoding *Hv*ERdj3B (w. SP) fused to the N-terminus of mYFP was co-transformed with an SP-mCherry-HDEL construct into barley epidermal cells using particle bombardment. The cells were observed by confocal laser scanning microscopy 48 h later. (c-f) Localization of *Hv*ERdj3B in *N. benthamiana* leaf epidermal cells using the split GFP system. *Hv*ERdj3B (w. SP)-GFP11 and (SP)-GFP1-10-HDEL reconstitute reticulate fluorescent GFP, while GFP11-*Hv*ERdj3B (w/o SP) and GFP1-10 reconstitute cytosolic and nuclear fluorescent GFP. Expression constructs were introduced using *Agrobacterium* infiltration, and epidermal cells were observed by laser scanning confocal microscopy 48 h later. Size bars, 20 μm.

To reveal *Hv*ERdj3B’s subcellular localization, it was expressed with its signal peptide and a C-terminal mYFP fusion in barley leaf epidermal cells. Laser scanning confocal microscopy indicated that *Hv*ERdj3B is an ER protein as the mYFP signal has a reticulate pattern, which overlaps with the (SP)-mCherry-HDEL ER marker (Figure 5b). This localization of *Hv*ERdj3B found in barley was confirmed in *N. benthamiana* (Figure S8). Next, we tested if *Hv*ERdj3B, as expected, is an ER luminal protein. Here we made use of a split GFP system different from BiFC. GFP is barrel-shaped and consists of 11 β-sheets (GFP1-11), which can be split into two fragments, β-sheets 1– 10 (GFP1-10) and β-sheet 11 (GFP11). The two parts of GFP have affinity for one-another, and when they co-localize in the same compartment, a fluorescent GFP complex will be assembled (Xie et al., 2017). Thus, when a *Hv*ERdj3B-GFP11 fusion protein was co-expressed with an SP-GFP1-10-HDEL construct that delivers and retains GFP1-10 in the ER lumen, perinuclear and reticulate GFP signals appeared (Figure 5c). No visible signal appeared when *Hv*ERdj3B-GFP11 was co-expressed with cytosolic GFP1-10 (Figure 5e). Furthermore, when GFP11-*Hv*ERdj3B was co-expressed with either (SP)-GFP1-10-HDEL or with cytosolic GFP1-10, GFP signal was absent or present in the cytosol and nucleus, respectively (Figure 5d,f). In summary, these results show that *Hv*ERdj3B is important for barley immunity to *Bh* and that it is located inside the ER lumen. This complements accumulating evidence of the role of ERQC and ERdj3B in plant immunity (Nekrasov et al., 2009; Tintor & Saijo, 2014).

### AVR_A1_ and BEC1016 translocate from the plant cytosol to the ER lumen

For interactions to occur between either AVR_A1_ or BEC1016 and *Hv*ERdj3B in barley epidermal cells attacked by *Bh*, the two effectors are required to be secreted from the fungus, when their signal peptides are removed, and to be taken up, either across the plant plasma membrane or the extrahaustorial membrane, into the plant cytosol. Secondly, in the plant cell, the effectors are required to pass the ER membrane into the ER lumen. Indirect evidence that AVR_A1_ translocates into the plant epidermal cell cytosol comes from the observation that it functions as an avirulence protein that binds to the cytosolic NLR-protein, MLA1, and triggers a hypersensitive response (HR) (Lu et al., 2016). The fact that the number of haustoria is increased when BEC1016 is expressed in the epidermal cell cytosol after particle bombardment, suggests that BEC1016 also functions inside the plant cell (Figure 1a). To inquire whether AVR_A1_ and BEC1016 can translocate across the ER membrane, we started by over-expressing AVR_A1_-GFP and BEC1016-GFP (both w/o SP) in *N. benthamiana* epidermal cells. Here they were both localized to the cytosol and nucleus together with free mCherry, and no obvious ER signal could be distinguished as there apparently was a complete overlap between the GFP and mCherry fluorescent signals (Figure 6). Expression in barley epidermal cells also suggested the effectors mainly to be cytosolic (Figure S9).

**Figure 6.**
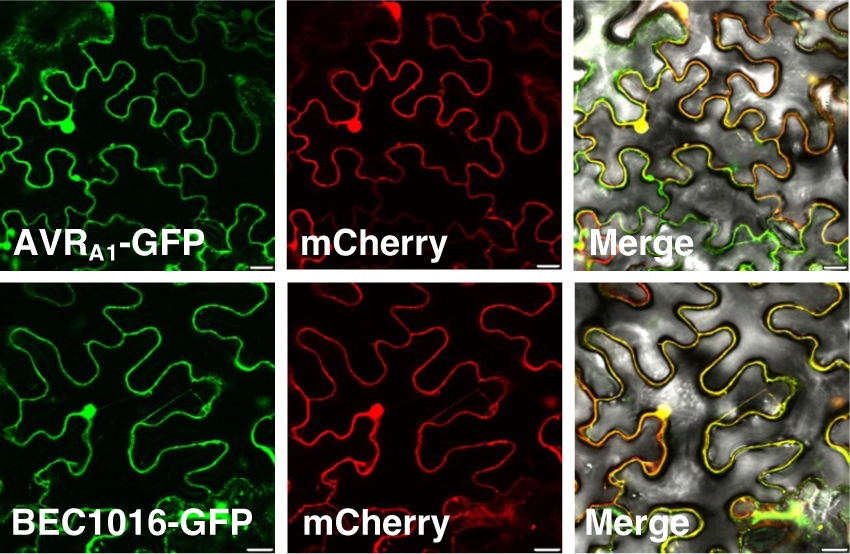
AVR_A1_ and BEC1016 localized in the cytosol and nucleus of *N. benthamiana* epidermal cells. Constructs for expression of effectors (w/o SP) were introduced using *Agrobacterium* infiltration, and epidermal cells were observed by laser scanning confocal microscopy 48 h later. Size bars, 20 μm.

However, to interact with *Hv*ERdj3B, the two effectors will have to enter the plant ER. To test this more rigorously, we again used our split GFP system (Xie et al., 2017). We generated constructs for fusing GFP11 to both the N and C-terminus of AVR_A1_, BEC1016 and CSEP0105, all w/o SP. CSEP0105, which interacts with cytosolic small heat shock proteins (Ahmed et al., 2015), was used as a negative control. These were then co-expressed with constructs for cytosolic GFP1-10 and ER-luminal (SP)-GFP1-10-HDEL in all combinations (Figure 7). GFP11 fused to the N and C-terminal of CNX^TM^, the transmembrane domain of the ER membrane protein, calnexin 1 (Xie et al., 2017), was used as reference (Figure 7a,b,i,j). All three GFP11-effector fusions combined with GFP1-10 resulted in cytosolic and nuclear GFP signal (Figure 7c-h). Yet, when combined with (SP)-GFP1-10-HDEL, removal of the cytosolic and nuclear GFP signal visualized distinct reticulate and perinuclear GFP signals in case of AVR_A1_ and BEC1016 (Figure 7k-n, S10). Since this was not seen for CSEP0105 (Figure 7o,p) and because the ER-localized GFP11-CNX^TM^/GFP1-10-HDEL combination also gave reticulate network signal (Figure 7i), the data indicate that a fraction of the cytosolic AVR_A1_ and BEC1016 specifically translocate into the ER lumen post-translationally and independently of their signal peptides.

**Figure 7.**
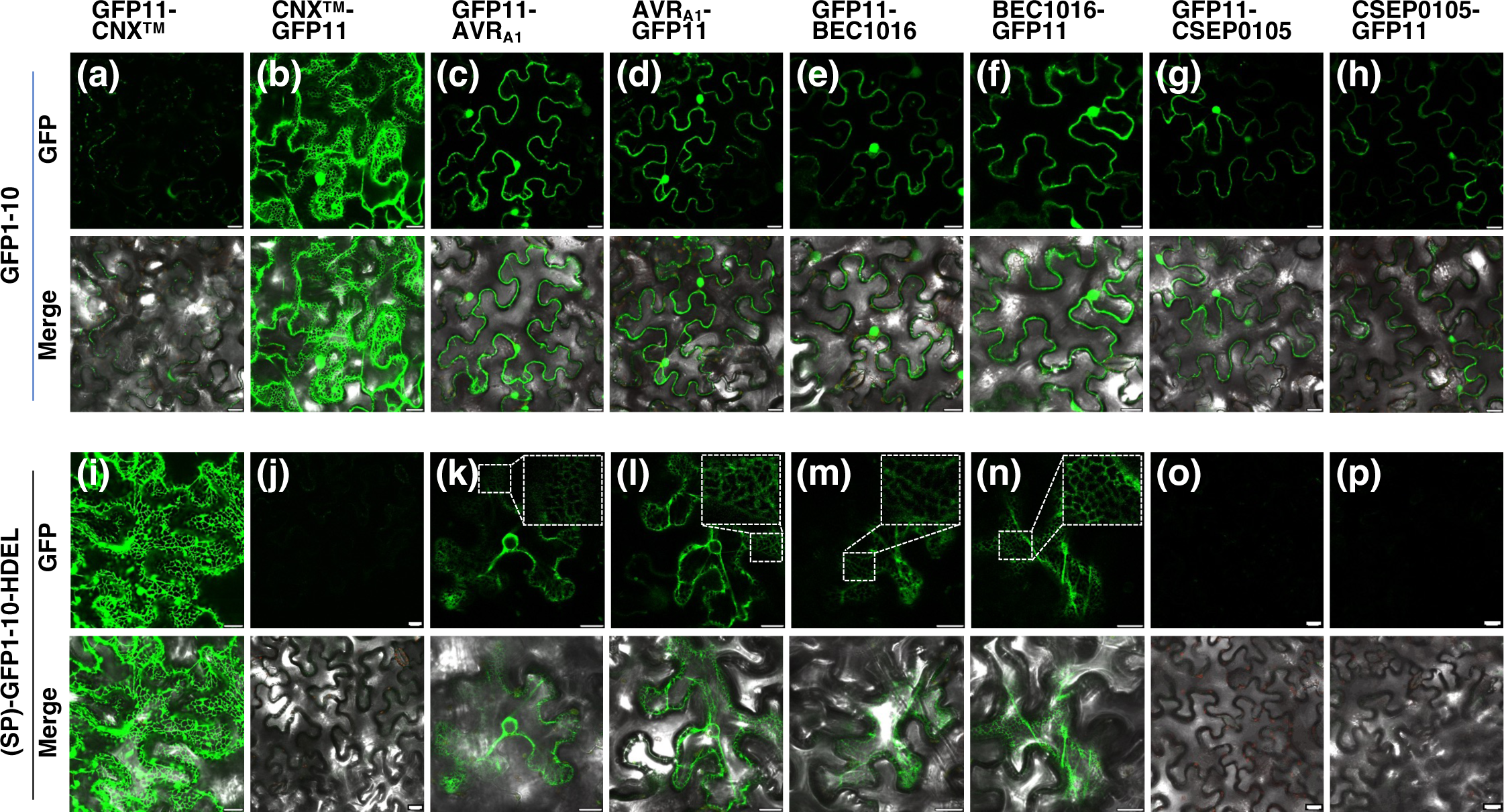
AVR_A1_ and BEC1016 are partially translocated into the ER lumen. The GFP1-10/GFP11 self-constituting fluorescence protein demonstrates cytosolic and ER-luminal protein localizations *N. benthamiana* epidermal cells. (a,b) and (i,j) GFP11-CNX^TM^ and CNX^TM^-GFP11 confirm (SP)-GFP1-10-HDEL to be ER-luminal and GFP1-10 to be cytosolic, respectively. (c-h) and (k-p) AVR_A1_, BEC1016 and CSEP0105 (all w/o SP) are cytosolic while AVR_A1_ and BEC1016 (both w/o SP) are also ER-lumunal, indicated by their reticulate and perinuclear signals (see also **Fig. S10**). Expression constructs were introduced using *Agrobacterium* infiltration, and epidermal cells were observed by laser scanning confocal microscopy 48 h later. Size bars, 20 μm.

### Silencing of *Hv*ERdj3B and expression of AVR_A1_ and BEC1016 affect ER trafficking

Having observed an importance of AVR_A1_, BEC1016 and *Hv*ERdj3B in immunity and localization of these three proteins in the plant ER lumen, we were prompted to test whether they can affect ER function. For this we made use of the vacuolar marker, RFP-AFVY, expressed with an N-terminal signal peptide. The C-terminal AFVY amino acid sequence, that guides the protein to the vacuole, is derived from the vacuolar storage protein, phaseolin (Hunter et al., 2007). When the (SP)-RFP-AFVY construct was expressed together with the empty-vector control, a clear vacuolar localization was observed (Figure 8). However, when co-expressed with AVR_A1_ or BEC1016, the RFP signal was also detected around the nucleus and to some extent in reticular structures, which are signs of (SP)-RFP-AFVY being retained in the ER. The same signal pattern was observed when *HvERdj3B* was silenced using the RNAi-1 hairpin construct, also used in Fig. 5A above (Figure 8).

**Figure 8.**
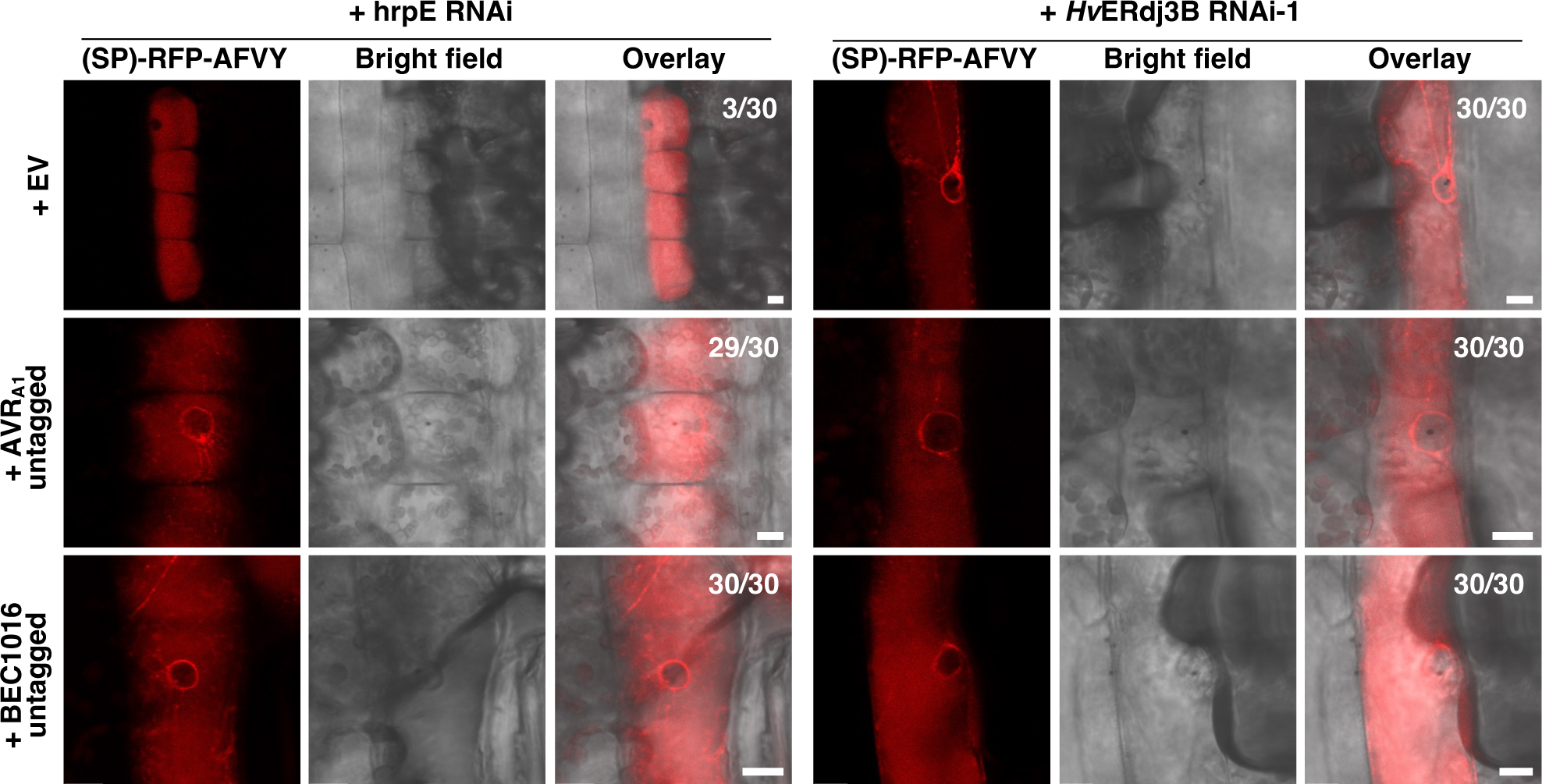
Overexpression of AVR_1A_ and BEC1016 or RNAi of the target *Hv*ERdjB3 cause ER retention of a vacoular marker. The vacuolar marker construct, p35S::SP-RFP-AFVY, was co-expressed with either the pUbi::GW empty vector (EV), pUbi::AVR_A1_ or pUbi::BEC1016 (both w/o SP), along with either the control RNAi construct (hrpE-RNAi) or an RNAi construct for *Hv*ERdj3B, in barley epidermal cells upon particle bombardment. The vacuolar marker, (SP)-RFP-AFVY, was partially mis-localized in an ER-like structure after overexpression of AVR_A1_ and BEC1016 or after RNA-interference of *Hv*ERdj3B. This was evident by its presence in a perinuclear ring and occasionally in reticular structures. Counts indicate occurrence of mis-localization of the vacuolar marker in 30 cells. Scale bars, 10 µm.

When expression of the effectors and silencing of *HvERdj3B* was combined, a similar pattern of the RFP signal was again seen. This shared cellular phenotype, induced by expression of the effectors and silencing of *HvERdj3B*, agrees with hampering of the ERQC and with this J-domain protein being targeted by the effectors in the ER lumen.

## DISCUSSION

Powdery mildew fungi are obligate biotrophic pathogens that feed on living plant tissues for nutrient uptake via haustoria. To suppress host defenses and promote colonization, these pathogens deliver a large repertoire of effectors into host cells (Frantzeskakis et al., 2018). Therefore, identifying the host targets of these effectors and their function will lead to detailed understanding of how infection is established and maintained. We focused on *Bh* AVR_A1_ and BEC1016, as examples of avirulence proteins and proteins having a detectable contribution to fungal virulence, respectively (Lu et al., 2016; Pliego et al., 2013). Here we found that bacterial T3SS-based delivery of these effectors reduced defense-associated callose deposition and that over-expression of BEC1016 enhanced the development of *Bh* haustoria. Through Y2H-NGIS, these effectors were found to target the same host protein, *Hv*ERdj3B, which we further substantiated with binary Y2H as well as *in planta* BiFC and co-IP assays. Silencing of the *HvERdj3B* gene in barley led to increased formation of *Bh* haustoria, indicating that this ER protein is involved in the plant’s immunity against fungal penetration. This, and the observation that AVR_A1_ and BEC1016 can enter the ER, and that expression of them as well as silencing of *Hv*ERdj3B cause arrest of a vacuolar marker in the ER, supports the immunity-suppressing function of these effectors to be mediated through targeting of this J-domain protein.

It is well-documented that ER stress responses and ERQC contribute to plant immunity. Our data are consistent with a previous study showing that a T-DNA line mutated in *HvERdj3B*’s closest homologue in *Arabidopsis*, *AtERdj3B*, is more susceptible to bacterial pathogens (Nekrasov et al., 2009). Here it was reported that *At*ERdj3B is involved in PTI. In the ER, it forms a complex with SDF2 and the Hsp70 BiP1, which are required for proper trafficking and function of the plasma membrane leucine-rich-repeat receptor kinases, EFR. *Hv*ERdj3B may have a similar function and may be involved in proper functioning of PRRs in barley.

Besides being required for processing of PRRs, ERQC is required for secretion of antimicrobial proteins and linked to pathogen-induced programmed cell death (Li et al., 2009; Kørner et al., 2015; Moreno et al., 2012; Qiang et al., 2012). In addition, a recent study showed that the ER proteins, UBAC2 and PICC, interact with each other and are required for proper delivery of callose synthase to the plasma membrane and for flagellin-triggered callose deposition (Wang et al., 2019). As studies in Arabidopsis have demonstrated that callose is important for immunity towards penetration by powdery mildew fungi (Voigt, 2016), it will be interesting to see if future studies can demonstrate that the ERdj3B/BiP complex is required for UBAC2/PICC-dependent callose deposition. Therefore, even if ERdj3B is essential for maturation of certain PRRs in the ER, there might be additional ways by which it contributes to plant immunity and thus how *Bh* uses AVR_A1_ and BEC1016 to aid the penetration process in barley.

Our immediate localization study suggested both effectors to be cytosolic and nuclear, while *Hv*ERdj3B was found to be ER-luminal. Therefore, we used the split GFP system to demonstrate that these effectors are also translocated into the ER post-translationally. There are other studies showing that pathogen effectors enter the ER to facilitate infection by targeting ER-localized host proteins. For instance, the RXLR effector PcAvr3a12 from the *Arabidopsis* oomycete pathogen, *Phytophthora capsici*, can post-translationally enter the ER and target the ER-localized host PPIase, FKBP15-2, which is involved in ER stress-sensing and required for ER stress-mediated plant immunity (Fan et al., 2018). Furthermore, it has been reported that an essential effector, PsAvh262, secreted by the soybean pathogen, *P. sojae*, translocates from the host cytosol into the ER to stabilize BiPs, thereby suppressing ER stress-triggered cell death and facilitating infection (Jing et al., 2016). In both cases, it was shown by BiFC that effector-target interactions occur inside the ER. Yet another recent study showed that a nematode effector (CLE) is translocated from the plant cell cytosol to the ER, and then secreted as an extracellular receptor ligand (Wang et al., 2021). Here post-translational ER uptake was demonstrated using the GFP1-10/11 system, which we also used. Together these observations suggest that these effectors hijack one or more signal peptide-independent plant ER uptake mechanisms.

AVR_A1_ and BEC1016 both interact with *Hv*ERdj3B and both enter the ER post-translationally and signal peptide-independently. They have very diverse amino acid sequences, but appear derived from an ancient RNAse and may potentially be structurally related (Cao et al., 2023; Pedersen et al., 2012), which may explain their shared functions. Interestingly, the ERQC machinery is known to engage in the post-translational ER protein uptake through the Sec61 translocon complex (Hassdenteufel et al., 2019; Zimmermann et al., 2011). Thus, we consider that interactions of AVR_A1_ and BEC1016 with *Hv*ERdj3B may facilitate one-way movement through this translocon into the ER lumen. Previously, we found that silencing of the barley Sec61 β-and γ-subunits both reduce the plant’s susceptibility to *Bh* (Xu et al., 2015; Zhang et al., 2013). These findings were at the time difficult to explain as preventing ER uptake of immunity-associated proteins, such as PR-proteins and PM-bound PRR, should increase the plant susceptibility. However, our present finding of a susceptibility promoting role of AVR_A1_ and BEC1016 (Figure 1), which both enter the ER, may make the consequences of Sec61 silencing (Xu et al., 2015; Zhang et al., 2013) more meaningful.

Often effectors are recognized indirectly by NLRs after they have impacted effector targets, which are guarded and associated with the NLRs (Carter et al., 2019; Ngou et al., 2022; Thordal-Christensen, 2020). However, an effector target that contributes to immunity, like *Hv*ERdj3B, inside the ER may not be guarded by an NLR, as to date, such receptors have not been localized in this compartment. Instead, this favors a requirement for direct effector monitoring by NLRs in the cytosol, which agrees with the direct recognition and interaction of AVR_A1_ and the NLR, MLA1 (Lu et al., 2016). One may speculate whether targeting an ER luminal host protein thus has provided an evolutionary benefit to the pathogen, since it may be more demanding for the plant to invent specific recognition of each effector, rather than guarding proteins targeted by more effectors potentially from different pathogens.

## EXPERIMENTAL PROCEDURES

### Plant and Fungal Materials

Seedlings of barley (*Hordeum vulgare* L.) cv. ‘Golden Promise’ were grown at 16 h light (150 μE s^−1^ m^−2^, 20°C) / 8 h of darkness (20°C) to be used for inoculations, gene amplifications, TIGS, overexpression, and callose deposition assays after effector-delivery using EtHAn. *Bh* isolate C15 (*AVR_a1_*) was propagated on barley P-02 plants in a cycle of 1 week. Four to six-week-old *N. benthamiana* plants were used for BiFC and subcellular localization studies after Agrobacterium-mediated leaf cell transformation.

### Gateway Plasmid Construction

<u>C</u>oding <u>D</u>NA <u>s</u>equences (CDS) for the effectors, AVR_A1_ and BEC1016, w/o their signal peptides, were amplified using primer pairs listed in Table S2. PCR was performed on cDNA from barley leaves infected with *Bh*, and the fragments were cloned into pENTR/D-TOPO® vectors (Invitrogen), with or without stop codons. The CDS for the full-length barley *Hv*ERdj3B (HORVU1Hr1G022990; GenBank ID: AK376215.1 and KAE8793706) with stop codon was synthesized by TWIST Bioscience (San Francisco, CA, USA). Fragments of *Hv*ERdj3B were amplified from the synthesized clone using primers listed Table S2 and cloned into pENTR/D-TOPO. Subsequently, the entry clone inserts were transferred to destination vectors using Gateway LR cloning reactions (Invitrogen). For over-expression constructs, the CDSs for *Hv*ERdj3B (with signal peptide), AVR_A1_ and BEC1016 (w/o signal peptides) were transferred into pUbi-mYFP-Gateway-Nos, pUbi-Gateway-mYFP-Nos, pUbi-Gateway-Nos destination vectors (Kwaaitaal et al., 2010) and p2WFHB-Gateway-GFP (Karimi et al., 2002). The vacuolar marker SP-RFP-AFVY construct, where the AFVY vacuolar targeting signal is derived from phaseolin, was amplified from ‘sp-RFP-AFVY’ (Hunter et. al, 2007). The SP-mCherry-HDEL construct was from Nelson et al. (2007). RNAi constructs were generated in the 35S promoter-driven hairpin destination vector, pIPKTA30N (Douchkov et al., 2005). The *Hv*ERdj3B-RNAi-1 and −2 constructs contain *Hv*ERdj3B CDS fragments from positions 283 to 579 and 532 to 879, respectively. These sequences were predicted using the si-Fi21 open-source software (Lück et al., 2019). None of them were found to have off-targets. For co-IP, the CDS for AVR_A1_ and BEC1016 (w/o signal peptide) were transferred into destination vector, pEarleyGate201, which is a Gateway-compatible vector encoding an N-terminal HA tag (Earley et al., 2006). The CDS of HvERdj3B (w/o signal peptide) was transferred into destination vector, pK7FWG2, a Gateway-compatible vector encoding a C-terminal GFP tag (Karimi et al., 2002). All the constructs were sequenced for confirmation.

### Barley Single Cell Transient-Induced Gene Silencing and Over-Expression

Barley leaf epidermal cell transformation was obtained after particle bombardment as described by Douchkov et al. (2005) and Nowara et al. (2010), using the biolistic PDS-1000/He Particle Delivery System from Bio-Rad. For each bombardment, six detached 1^st^ leaves of one-week-old Golden Promise barley plants were used. The particle coating was performed using 7 μg of DNA for each construct together with 2.4 mg of gold, 1 μg μl^−1^ of protamine (Sivamani et al., 2009) and 0.625 M CaCl_2_ (Rasco-Gaunt et al., 1999). For the bombardments, a hepta adapter and rupture discs bursting at a helium pressure of 1,100 psi were used. After bombardment, the leaves were transferred onto 1% phytoagar petri dishes containing 40 mg ml^−1^ benzimidazole. For effector over-expression and TIGS studies, the constructs were co-transformed with a GUS reporter gene construct driven by the pUbi promoter into the epidermal cells. Two days later, the leaves were inoculated with *Bh*, and after another 2 days, they were stained for GUS activity by vacuum-infiltrating and incubating in 2 mM X-Gluc, 100 mM sodium phosphate, 100 mM EDTA, 1.4 mM potassium ferricyanine, 1.4 mM potassium ferrocyanine, 0.1% Triton X-100 at 37°C overnight.

Transformed (blue) cells were assessed microscopically for the presence of haustoria as an indication of fungal infection. The haustorium formation rate was calculated as the number of blue cells with haustoria divided by the total number of blue cells. A mYFP and the empty vector pIPKTA30N were used as negative controls and the Mlo RNAi (pIPKTA36) (Douchkov et al., 2005) construct was used as positive control. The effect of constructs was analysed by a logistic regression model with random effects, assuming a binominal distribution, with construct as fixed effect and the experiment × construct × leaf interaction as random effect. The analyses were performed in PC-SAS (release 9.4, SAS Institute, Cary, NC).

### *A. tumefaciens* infiltration-mediated transformation of *N. benthamiana* leaf cells

T-DNA construct were introduced into *A. tumefaciens* strain GV3101 by electroporation. The transformed bacterial cells were grown on LB agar plates supplemented with Rifampicin, Spectinomycin and Gentamycin antibiotics. For leaf cell transformation, over-night liquid cultures (28°C) of recombinant *A. tumefaciens* were harvested by centrifugation and resuspended in 10 mM MgCl_2_, 10 mM MES and 200 μM acetosyringone to OD_600_=0.6. A strain of *A. tumefaciens* with a construct expressing the p19 silencing suppressor (Brioudes et al., 2022) was managed the same way. The resuspended *A. tumefaciens* transformants, including the one with the P19 construct, were mixed in equal ratios and infiltrated into leaves of four to six-week-old *N. benthamiana*.

### Yeast Two-Hybrid, Next-Generation Interaction Screen

The *Bh* effectors AVR_A1_ and BEC1016 were used as baits in Y2H-NGIS to mine for novel PPI (Elmore et al., 2023; Velásquez-Zapata et al., 2023a). Bait sequences (w/o signal peptides) were fused with the GAL4 transcription factor binding domain (GAL4-BD) in the p97-BD, Leu2p vector and transformed into *Saccharomyces cerevisiae* strain Y8930 (Dreze et al., 2010). For the prey library, first seedling leaves from an infection time course of the resistant barley line CI 16151 (*Mla6*) and four fast-neutron-derived immune mutants were sampled from a split-plot design at 0, 16, 20, 24, 32, and 48 hours after inoculation (HAI) with *Bh* isolate 5874 (*AVR_a1_, AVR_a3_, AVR_a6_, AVR_a12_*) (Chapman et al., 2021; Surana, 2017; Velásquez-Zapata et al., 2022). mRNA isolated from the 90 experimental units (5 genotypes × 6 time points × 3 biological replications) was used for RNA-Sequencing (data at https://www.ncbi.nlm.nih.gov/geo/query/acc.cgi?acc=GSE101304), small RNA sequencing (Hunt et al. 2019), and was also pooled to prepare a three-frame cDNA prey library in the Gateway-compatible ARS4/CEN6 GAL4 activation domain (AD) vector, p86-AD, Trp1p (Dreze et al. 2010; Velásquez-Zapata et al., 2021; Yu et al. 2015). After transforming the library into *S. cerevisiae* strain Y8800, 1.1 x 10^7^ primary clones were mated with each bait clone and subsequently cultured in three independent replicates under two conditions (selection and non-selection). At the end of each culture, prey plasmids were isolated, and prey cDNA fragments were amplified by low-cycle PCR to maintain the bait/prey ratios resulting from the batch culture mating. Amplicon libraries were sequenced using the Illumina HiSeq 2500 platform at the Iowa State University DNA facility, collecting 5–10 million reads per sample. Sequence output from NGIS screens confirmed that the prey library contained 78.4% of annotated genes in the barley Morex V3 assembly (Mascher et al., 2021), and nearly 99% of the expressed genes from our CI 16151 transcriptome (NCBI-GEO GSE101304). Y2H-NGIS data from these experiments were analyzed using the NGPINT and Y2H-SCORES software (Banerjee et al., 2021; Velásquez-Zapata et al., 2021) to reconstruct the prey fragments and rank them as interactors for each bait. Outputs from these pipelines allowed us to identify interacting prey fragments, their frame with the Gal4-AD as well as enrichment and specificity scores that assess their properties as Y2H interactors.

### Binary Yeast 2-Hybrid Assays

Using the output from the Y2H-NGIS, we designed primers to re-clone identified prey fragments into p86-AD to confirm interactions using binary Y2H (Dreze et al., 2010) under three levels of selective media: Diploid selection (SC-LW), and specific selection (SC-LWH + 0.1 mM 3-AT) using three dilutions (10^0^, 10^-1^, 10^-2^) as shown in Figure S6. After the first confirmation of *Hv*ERdj3B as prey, new AVR_A1_, BEC1016 and *Hv*ERdj3B constructs were made in the destination vectors pDEST-AS2-1 (GAL4-BD) and pDEST-ACT2-1 (GAL4-AD) (Robertson, 2004). These constructs were transformed into the haploid yeast strains, Y189 and Y190, respectively. Subsequent matings, selections and *LacZ* reporter assays were made according to the Matchmaker Gold Yeast Two-Hybrid System user manual (Clontech; Mountain View, Ca, USA).

### *In Planta* Protein-Protein Interaction Studies

For the BiFC assay, the CDS for full-length barley *Hv*ERdj3B and both effectors, w/o signal peptides, and with and w/o stop codons, were transferred to 35S promoter-driven BiFC binary destination vectors (Kamigaki et al. (2016) as N- and C-terminal fusions of nGFP (aa 1 to 174) and cGFP (aa 175 to 239). Two-by-two combinations of constructs were transformed into *N. benthamiana* leaves by *A. tumefaciens* infiltration. Two days after infiltration, the fluorescence signals of all eight possible AVR_A1_/*Hv*ERdj3B combinations and of all eight possible BEC1016/*Hv*ERdj3B combinations were evaluated by a laser scanning confocal microscopy.

For the co-IP assay, *N. benthamiana* was agro-infiltrated with constructs for expression of *Hv*ERdj3B-GFP in combination with either HA-Avr_A1_ or HA-BEC1016, all without signal peptides. Two days later, co-IP was performed according to Gruner et al. (2021) using 15 μl α-GFP-magnetic beads (Chromotek). Anti-GFP antibody (sc9966), anti-HA antibody (sc7392 HRP) and secondary antibody m-IgGκ BP-HRP (sc-516102) were purchased from Santa Cruz Biotechnology.

### Localization of Effectors in the ER

The GFP1-10/GFP11 split GFP system (Xie et al., 2017) was used to document presence of effectors in the ER lumen. CDS for the effectors AVR_A1_ and BEC1016, w/o their signal peptides, were transferred to the T-DNA Gateway destination vectors, pGFP11-GW and pGW-GFP11, to fuse them to GFP11. T-DNA constructs for GFP11-CNX^TM^ and CNX^TM^-GFP11 were from Xie et al. (2017). These were co-expressed with constructs for cytosolic GFP1-10 and ER-luminal SP-GFP1-10-HDEL from Xie et al. (2017).

### Confocal Microscopy

The microscopy was performed using a Leica SP5-X laser scanning microscope at the Center for Advanced Biomaging (CAB) at the University of Copenhagen. To improve the subcellular localisation, the leaves were mounted with Perfluorodecalin (Alfa Aesar A18288) and imaged with a 20x water immersion lens. GFP was excited at 488 nm and the emission from 514 to 540 nm was collected. mYFP was excited at 514 nm and the emission from 527 to 586 nm was collected. mCherry was excited at 587 nm and the emission from 595 to 650 nm was collected. To restrict bleed-through, all imaging was done using sequential scan mode.

### Callose Deposition Assay

CDSs for the GUS reporter gene, effectors AVR_A1_ and BEC1016, w/o signal peptides and with stop codons were transferred to the AvrRPS4 promoter-driven destination vector, pEDV6 (Fabro et al., 2011), and transformed into *P. fluorescens* EtHAn (Thomas et al., 2009) by electroporation. Transformed EtHAn strains were grown over-night in King’s B medium containing ampicillin, chloramphenicol, tetracycline and gentamicin at 28°C. The bacteria were harvested by centrifugation and resuspended in 10 mM MgCl_2_ to a final OD_600_=0.3 and infiltrated into leaves of eight-day-old barley. Twenty-four hours later, the callose response was assayed in the infiltrated leaf areas by aniline blue fluorescence using a Nikon ECLIPSE Ni-U fluorescence microscope. For one repeat, ten 1110 μm x 740 μm (5184 × 3456 pixels) images from random sites at one leaf were used to quantify the number of callose deposits and the accumulated area of callose deposits using the Fiji open-source platform (Jin & Mackey, 2017; Schindelin et al., 2012). The outcomes of effectors were calculated relative to GUS as control and analysis of the callose deposition data was performed for the total number and area of the callose deposits independently. We performed a repeated measure ANOVA (Keselman et al., 2001) to test the effect of the treatments considering the repeated measures by leaf. Significant treatment effects were further explored with a paired-sample t-test and the *P*-values were adjusted using the method of Benjamini & Hochberg (1995). The GUS assay was performed as above. Growth of each EtHAn strain in barley leaves was quantified by extracting bacteria from homogenized leaves at 0 and 1 <u>d p</u>ost infiltration (dpi). The colony forming units assessed from the 1 dpi samples was normalizing to the average of those obtained from the 0 dpi samples.

### Differential transcript accumulation

RNA-Seq data were extracted from an infection time course of barley CI 16151 and the derived fast-neutron mutant, *mla6*-m18982, at 0, 16, 20, 24, 32, and 48 hours after inoculation with *Bh* isolate 5874 (*AVR_a1_, AVR_a6_*; NCBI-GEO GSE101304) and analyzed as described in (Chapman et al., 2021; Velásquez-Zapata et al., 2022). Genes differentially expressed at an adjusted *P* of < 0.001 for barley and < 0.003 for *Bh* were considered significant.

## DATA AVAILABILITY

Infection-time-course RNA-Seq data are at NCBI-GEO under the acc. no. GSE101304 (https://www.ncbi.nlm.nih.gov/geo/query/acc.cgi?acc=GSE101304). R code and the ReadMe file for the NGPINT and Y2H-SCORES software used to identify *Bh* AVR_A1_ and BEC1016 interactors are provided at the GitHub pages (https://github.com/Wiselab2/NGPINT_V2; https://github.com/Wiselab2/Y2H-SCORES). Raw Y2H-Seq reads are at NCBI-GEO under acc. nos. GSE164954 (AVR_A1_/CSEP0008) and GSE166108 (BEC1016/CSEP0491). Otherwise, Y2H-NGIS scores are available in the supporting information.

## ACCESSION NUMBERS

GenBank acc. nos. for *AVR_A1_* (CSEP0008, BLGH_03023) and *BEC1016* (CSEP0491, BLGH_07006), are CCU81904.1 and CCU83284.1, respectively. *Hv*ERdjB3 is represented in GenBank as an mRNA-derived DNA sequence from Haruna Nijo (AK376215.1) and as a protein sequence from the genome sequence of Tibetan Hulless barley (KAE8793706).

## ACKNOWLEDGEMENTS

We thank our late and good friend, Dr. Patrick Schweizer (Leibniz Institute of Plant Genetics and Crop Plant Research, Gatersleben, Germany) for providing the RNAi vector and the *Mlo* RNAi construct, Dr. Masumi Robertson (Commonwealth Scientific and Individual Research Organization Plant Industry) for the Gateway Y2H vectors (pDEST-ACT2 and pDEST-AS2-1), Greg Fuerst (USDA-ARS at Iowa State University) for conducting the *Bh* time-course infection experiment and expert isolation of RNA for the 3-frame Y2H library, Associate Professor Shoi Mano for the BiFC-vectors, and Drs. Pietro Spanu (Imperial College, U.K.) and Ralph Panstruga (RWTH Aachen, Germany) for CSEP clones. Research supported in part by China Scholarship Council for PhD-student scholarship 201708340064 to ZZ, Novo Nordisk Foundation Challenge grant NNF19OC0056457 to HTC, Villum Fonden Experiment Programme grant 00028131 to HTC, Fulbright - Minciencias 2015 & Schlumberger Faculty for the Future fellowships to VVZ, USDA-ARS Postdoctoral Research Associateship and USDA-NIFA-ELI Postdoctoral Fellowship 2017-67012-26086 to JME, Oak Ridge Institute for Science and Education under U.S. Department of Energy contract number DE-SC0014664 to SB, and National Science Foundation - Plant Genome Research Program grant 13-39348, USDA-National Institute of Food and Agriculture grant 2020-67013-31184, USDA-Agricultural Research Service projects 3625-21000-067-00D & 5030-21220-068-000-D to RPW and Marie Skłodowska-Curie Actions Postdoctoral Fellowship project 101104193 to SD. The funders had no role in study design, data collection and analysis, decision to publish, or preparation of the manuscript. Mention of trade names or commercial products in this publication is solely for the purpose of providing specific information and does not imply recommendation or endorsement by the CSC, NNF, VF, USDA, NIFA, ARS, DOE, ORAU/ORISE, NSF or MSCA. USDA is an equal opportunity provider and employer.

## SUPPORTING INFORMATION

**Figure S1.** Attempted alignment of the amino acid sequences of AVR_A1_ (CSEP0008, BLGH_03023) and BEC1016 (CSEP0491, BLGH_07006).

**Figure S2.** Transcript accumulation on barley inoculated with *Bh* isolate 5874.

**Figure S3.** EtHAn-delivery of β-glucuronidase into barley leaf cells.

**Figure S4.** Growth of EtHAn in barley leaves is independent on effector construct.

**Figure S5.** Alignment of *Hv*ERdj3B with human *Hs*ERdj3 (NP_057390.1) and Arabidopsis *At*ERdj3B (At3g62600).

**Figure S6.** Binary Y2H used to validate interactions between AVR_A1_, BEC1016 and *Hv*ERdj3B

**Figure S7.** Bimolecular fluorescence complementation study of the interaction between AVR_A1_, BEC1016 and *Hv*ERdj3B.

**Figure S8.** Subcellular localization of *Hv*ERdj3B in *N. benthamiana* epidermal cells.

**Figure S9.** Localization of AVR_A1_ and BEC1016 in barley leaf epidermal cells.

**Figure S10.** BEC1016 can translocate into the ER.

**Table S1.** Summary Y2H-SCORES data for the AVR_A1_ and BEC1016 baits.

**Table S2.** Primer sequences for plasmid constructions.

## AUTHOR CONTRIBUTIONS

ZL: Transient effector OE and haustorial formation rate, extensive binary Y2H, protein subcellular localizations, contributed to writing.

VVZ: Y2H-NGIS, developed the Y2H-SCORES software to rank high-priority candidates, identified *Hv*ERdj3B, and validated by first binary Y2H, contributed to statistics, contributed to writing,

JME: Initiated the overall concept and experimental plan. Constructed the 3-frame barley/Blumeria Y2H library and developed Y2H-NGIS screen.

XL: Callose response to EtHAn infiltrations.

WX: Assisted and supervised subcellular localizations.

SD: Expression of the vacuolar marker with expression of AVR_A1_ and BEC1016 or RNAi of the target *Hv*ERdj3B

XT: Co-IP experiments of *Hv*ERdj3B with AVR_A1_ and BEC1016

SB: Developed the NGPINT software and analyzed Y2H-NGIS sequence count data. HJLJ: Contributed to statistics.

CP: Supervised and contributed to writing.

RPW: Initiated the overall concept and experimental plan, organized, supervised and contributed to writing.

HTC: Organized and supervised the work, wrote first draft, lead the writing and finalized the work

