## Supplementary material for "Powdery mildew effectors AVR_A1_ and BEC1016 target the ER J-domain protein *Hv*ERdj3B required for immunity in barley": Suppl. Figures

#### Figure S1

```
AVRA1      MKFYHSSFVSNLTIL-SLFLNINGAFSRTWQCPSGDILYEEDVYSHTNMNSFEHFPFKTIEDE
BEC1016    MKFFSSASTAALTCLLSLVPVAHG--EPYFKCTQNTIFTMAEL--QRDKVYFLETNAKH-GDP

AVRA1      VYENTSLSRGVVYKVLDVHPSSSSATTFAVFTSNK-----RVAVVYQQGYRNNRKFIEECT
BEC1016    LGPNGEAYPTHRY----SMIPIRGYTTTYLFQLVDEAPTFRVFEK-QSGEFYPCNLIEN
```

**Figure S1. Attempted alignment of the amino acid sequences of AVR$_{A1}$ (CSEP0008, BLGH_03023) and BEC1016 (CSEP0491, BLGH_07006).** Signal peptides underlined (black), predicted beta sheets underlined (green), and intron position in the encoding genes indicated by dashed line (orange).

#### Figure S2

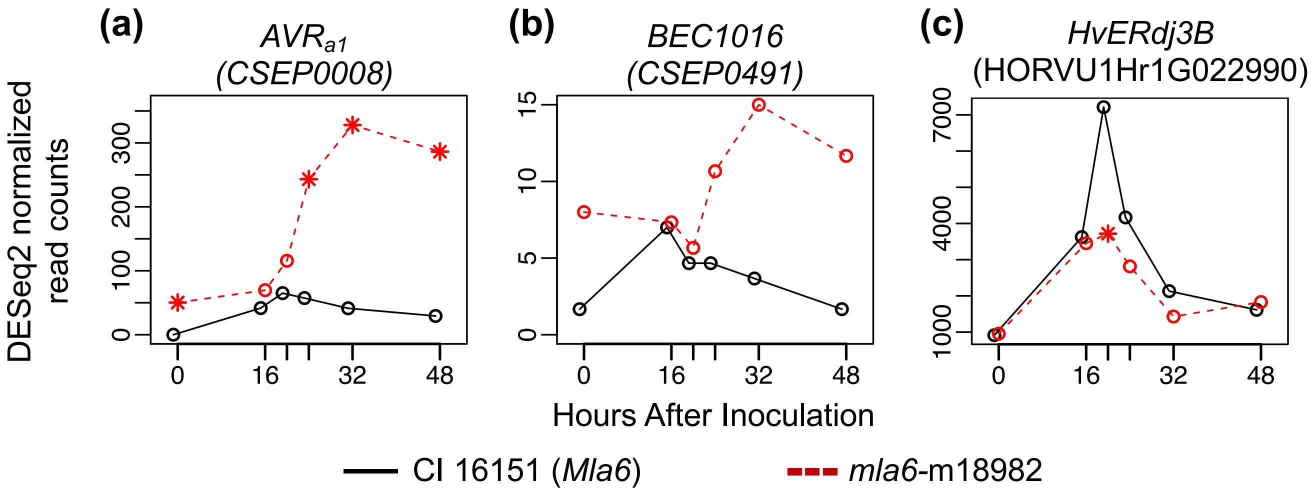


**Figure S2. Transcript accumulation on barley inoculated with *Bh* isolate 5874.** (a) *Bh* effector *AVRa1* (*CSEP0008*), (b) *Bh* effector *BEC1016* (*CSEP0491*) and (c) barley target *HvERdj3B* (HORVU1Hr1G022990) in wild-type progenitor CI 16151 (black solid lines) vs. fast-neutron derived susceptible mutant *mla6*-m18982 (dashed red lines). *, significant difference from CI 16151 at an adjusted $P < 0.001$ for barley genes. $P < 0.003$ for *Bh*.

### Figure S3

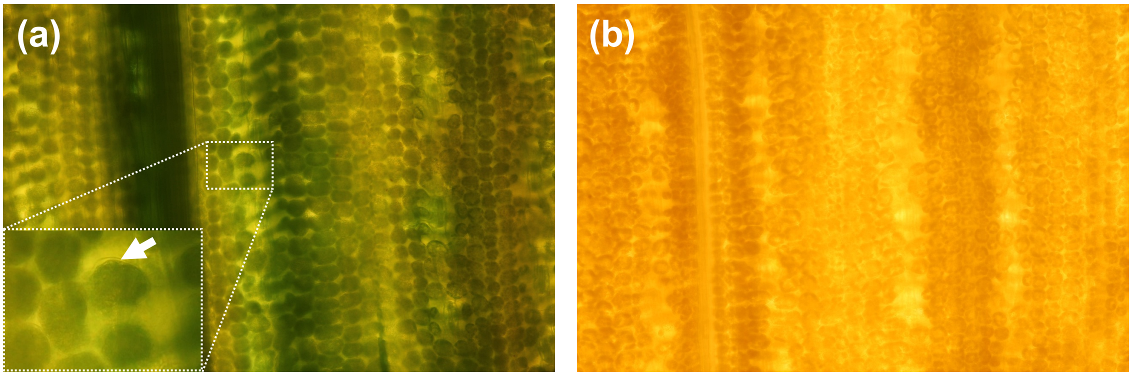


**Figure S3. EtHAn-delivery of β-glucuronidase into barley leaf cells.** (a) Leaf infiltrated with EtHAn transformed with pEDV6-GUS. Arrow, mesophyll cell wall. (b) non-infiltrated leaf. The two leaves were in parallel incubated in a GUS reaction buffer.

### Figure S4

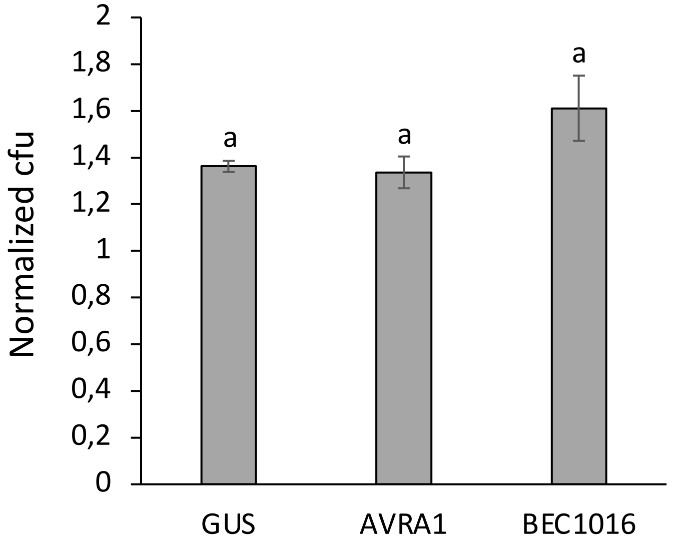


**Figure S4. Growth of EtHAn in barley leaves is independent on effector construct.** Colony-forming units (cfu) of Ethan strains extracted at 1 d after infiltration (dai) into 1$^{st}$ leaf of barley Golden Promise. The numbers were normalized to the average cfu obtained from at 0 dai. Error bars, SE. Letter (a) indicates no significant differences ($P$ < 0.05). Data were analyzed by a mixed effect-model analysis of variance. n=3.

### Figure S5

```
                    Signal peptide                                J-domain
HsERdj3     --MAPQNLSTFCL--LLLYLIGAVIAGRDFYKILGVPRSASIKDIKKAYRKLALQLHPDR     56
HvERdj3B    MAAPRRSGARLAAVLALLLHLAAVIEGKSFYDVLQVPKGASEDQIKRSYRKLALKYHPDK     60
AtERdj3B    MAI---RWSELCIVLFALSYAICVLAGKSYYDVLQVPKGASDEQIKRAYRKLALKYHPDK     57
                    : :.     *    .*: *:.:*.:* **:.** .:**::******: ***:

                                                               G/F-domain
HsERdj3     NPDDPQAQEKFQDLGAAYEVLSDSEKRKQYDTYGEEGLKDGH----------QSSHGDIF     106
HvERdj3B    NPDNEEATKRFAEINNAYEVLTDQEKRKVYDRYGEEGLKQFQ--GGRGGGGGMNMQDIF     118
AtERdj3B    NQGNEEATRKFAEINNAYEVLSDEEKREIYNKYGEEGLKQFSANGGRGGGGGMNMQDIF     117
            * .: :* .:* ::. *****:*.***: *: *******:              .  ***

                                          Substrate binding-domain
HsERdj3     SHFFGDFGFMFGGTPRQQDRNIPRGSDIIVDLEVTLEEVYAGNFVEVVRNKPVARQAPGK     166
HvERdj3B    SSFFGGGG----GGMEEEEEQIIKGDDVIVELDASLEDLYMGGSLKVWREKNIIKPAPGK     174
AtERdj3B    SSFFGGGS-------MEEEEKVVKGDDVIVELEATLEDLYMGGSMKVWREKNVIKPAPGK     170
            * ***. .        :::.:: :*.*:**:*:.:**::* *. ::* *:* : : ****
                                       142
                Cys-rich domain                       Substrate binding-domain
HsERdj3     RKCNCRQEMRTTQLGPGRFQMTQEVVCDECPNVKLVNEERTLEVEIEPGVRDGMEYPFIG     226
HvERdj3B    RRCNCRNEVYHRQIGPGMYQQMTEQVCDQCPNVKYVREGDFLTVDIEKGMQDGQEVSFFE     234
AtERdj3B    RKCNCRNEVYHRQIGPGMFQQMTEQVCDKCPNVKYEREGYFVTVDIEKGMKDGEEVSFYE     230
            *:****:*:   *:*** :*   * ***:*****  .*   : *:** *::** *  *

HsERdj3     EGEPHVDGEPGDLRFRIKVVKHPIFERRGDDLYTNVTISLVESLVGFEMDITHLDGHKVH     286
HvERdj3B    EGEPKIDGEPGDLKFRIRTAPHDRFRREGNDLHATVTISLLQALVGFEKNLKHLDNHLVQ     294
AtERdj3B    DGEPILDGDPGDLKFRIRTAPHARFRRDGNDLHMNVNITLVEALVGFEKSFKHLDDHEVD     290
            :*** :**:****:***:.. *  *.* *:**: .*.*:*::*****  .:.***.* *.
            Dimerization-domain
HsERdj3     ISRDKITRPGAKLWKKGEGLPNFDNNNIKGSLIITFDVDFPKEQLTEEAREGIKQLLKQG     346
HvERdj3B    IGSQGVTKPKEVRKFKGEGMPLHQ-SNKKGDLYVTFEVLFPKTLT-DDQKAKLKDVLA--     350
AtERdj3B    ISSKGITKPKEVKKFKGEGMPLHY-STKKGNLFVTFEVLFPSSLT-DDQKKKIKEVFA--     346
            *. . :*:*      ****:* .  .. **.* :**:* **.    :: :  :*:::

HsERdj3     SVQKVYNGLQGY      358
HvERdj3B    ------------      350
AtERdj3B    ------------      346
```

**Figure S5. Alignment of *Hv*ERdj3B with human *Hs*ERdj3 (NP_057390.1) and Arabidopsis *At*ERdj3B (At3g62600).** Alignment made using Clustal Omega. Domain structure defined by *Hs*ERdj3 according to Chen et al. (2017, EMBO J, 36: 2296). Valine 142, marking the N-terminus of the *Hv*ERdj3B protein encoded by the longest NGIS-Y2H reads, is highlighted (see also Fig. 2).

### Figure S6

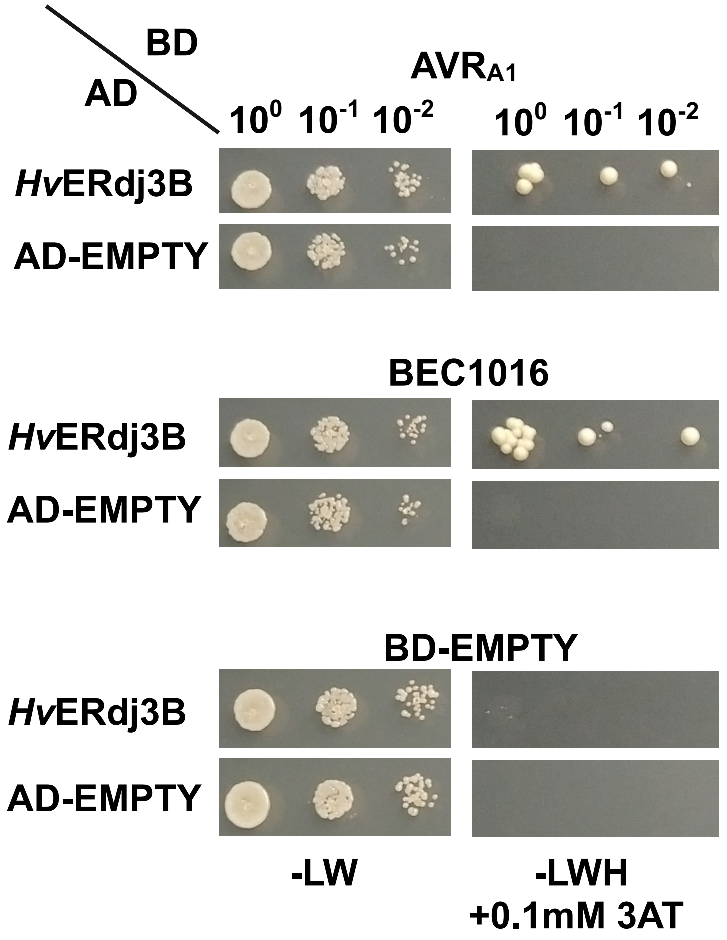


**Figure S6. Binary Y2H used to validate interactions between AVR$_{A1}$, BEC1016 and *Hv*ERdj3B.** Binary confirmation of the interactions between AVR$_{A1}$, BEC1016 and *Hv*ERdj3B (aa 132-350), using the binary Y2H system described by Dreze *et al.* (2010) using DO-LW media as diploid control and DO-LWH + 3AT as selection for positive interactions. Images taken 5 days after plating.

#### Figure S7

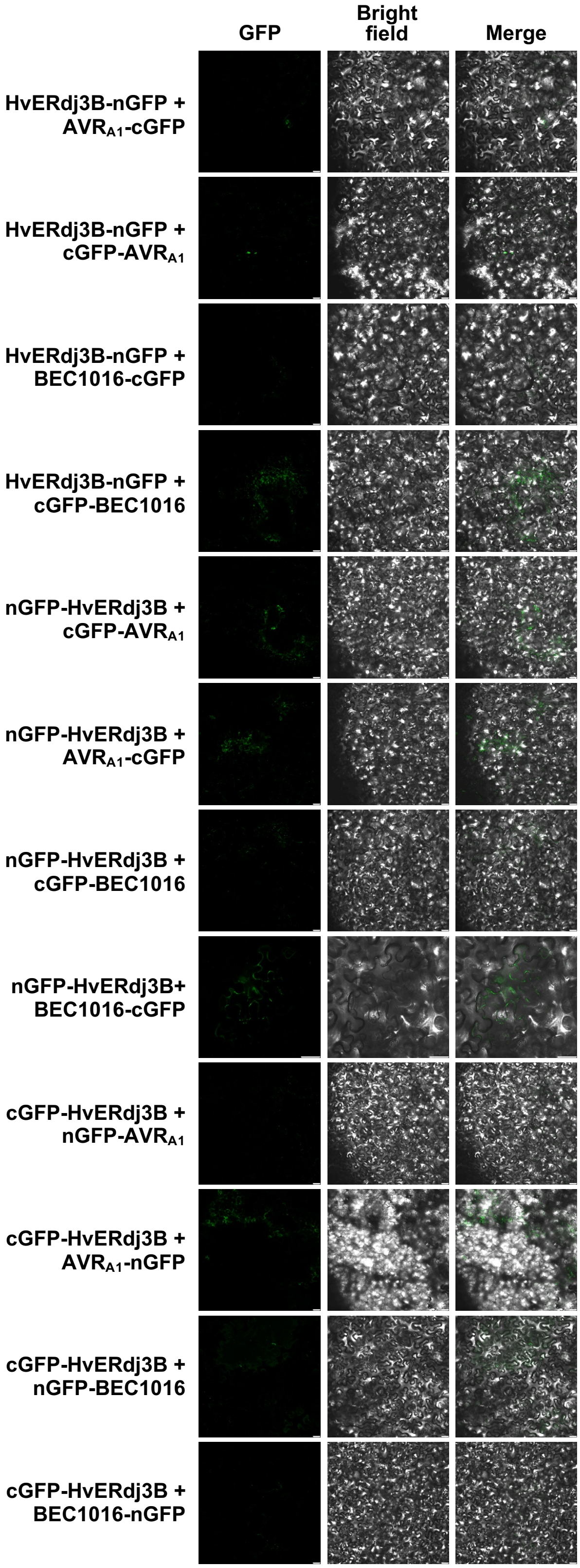


**Figure S7. Bimolecular fluorescence complementation study of the interaction between AVR$_{A1}$, BEC1016 and *Hv*ERdj3B.** These are the non-fluorescent combinations. The fluorescent combinations involving *Hv*ERdj3B-cGFP are shown in **Fig. 4.** See legend of **Fig. 4** for more details. Size bars, 40 μm.

### Figure S8

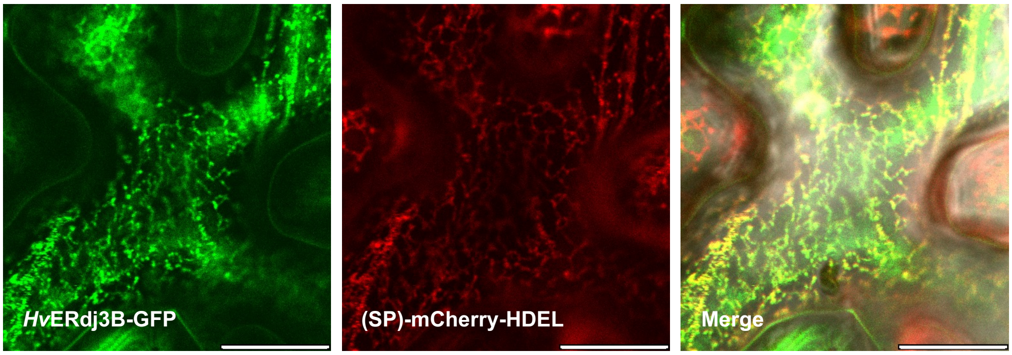


**Figure S8. Subcellular localization of *Hv*ERdj3B in *N. benthamiana* epidermal cells**. *Hv*ERdj3B (w. SP) gave reticular signal overlapping with the (SP)-mCherry-HDEL ER marker. Size bars, 20 μm.

### Figure S9

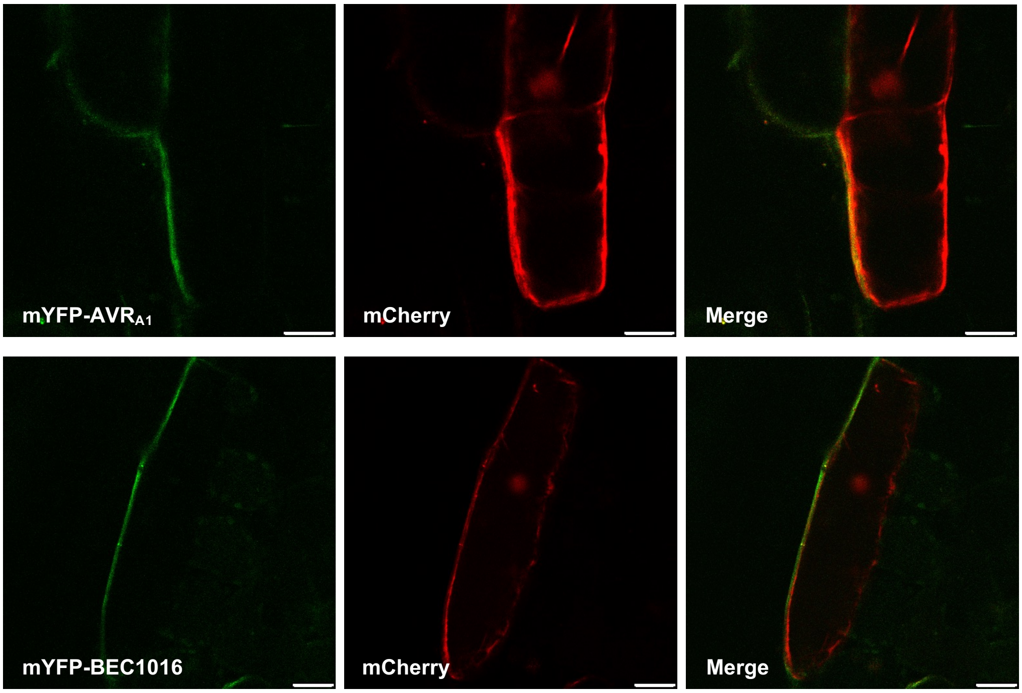


**Figure S9. Localization of AVR$_{A1}$ and BEC1016 in barley leaf epidermal cells.** Ubiquitin promoter-driven expression constructs encoding SP-free effectors (fused to the C-terminus of mYFP) were co-transformed with a free mCherry construct into barley epidermal cells using particle bombardment. The cells were observed by confocal laser scanning microscopy 48 h later. Size bars, 20 μm.

### Figure S10

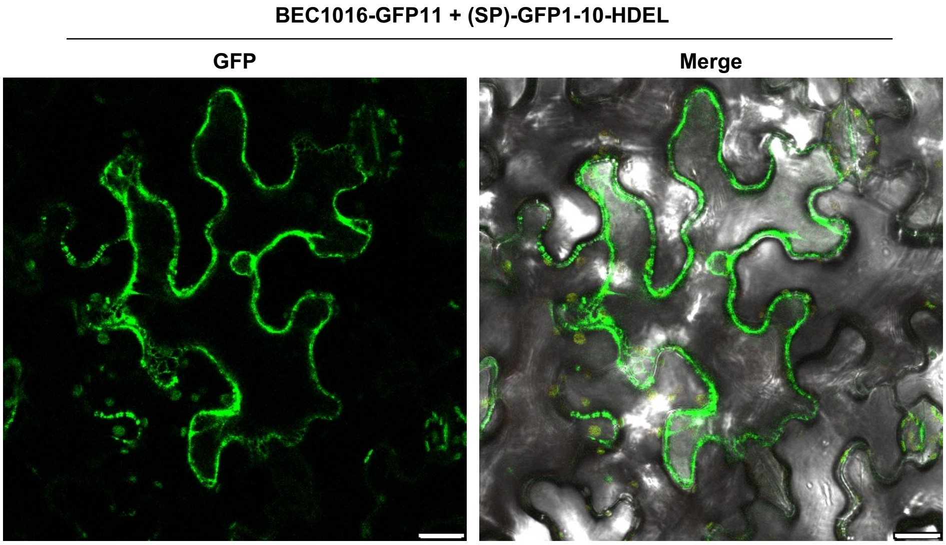


**Figure S10. BEC1016 can translocate into the ER.** Construct encoding BEC1016-GFP11 (w/o SP) and SP-GFP1-10-HDEL were co-transformed into *N. benthamiana* leaves using *Agrobacterium* infiltration, and epidermal cells were observed by laser scanning confocal microscopy 48 h later. The ER localization is indicated by the reticulate and perinuclear signals. See also **Fig. 7**. Size bars, 20 μm.
