## Supplementary material for "Powdery mildew effectors AVR_A1_ and BEC1016 target the ER J-domain protein *Hv*ERdj3B required for immunity in barley": Suppl. Table 2

**Table S2. Primer sequences for plasmid constructions** (5´ to 3´)**.**

| **Primers for the bait and prey constructs in Y2H and BiFC** | |
| --- | --- |
| HvERdj3B-FL_F | CACCATGGCCGCCCCCAGA |
| HvERdj3B-FL_R no stop | TGCAAGAACATCCTTCAGCTT |
| HvERdj3B-FL_R with stop | TTATGCAAGAACATCCTTCAGCTT |
| HvERdj3B_Fw without SP | CACCATG AAGAGCTTCTACGACGTGCTG |
| HvERdj3B_Rw with stop | TTATGCAAGAACATCCTTCAGCTT |
| HvERdj3B-C-term_F | CACCATGGTGATTGTTGAACTGGATGCTT |
| HvERdj3B-C-term_R with stop | TTATGAAACCTCCTGTCCATCTTG |
| CSEP0008_F without SP | CACCATGcgaacgtggcaatgccca |
| CSEP0008_R no stop | ggtgcattcttcaatgaattt |
| CSEP0008_R with stop | TTAGGTGCATTCTTCAATGAATTT |
| CSEP0491_F without SP | CACCATGgaaccatattttaaatgtactcaaaac |
| CSEP0491_R no stop | attttctatcaaattgcatggata |
| CSEP0491_R with stop | TTAATTTTCTATCAAATTGCATGGATA |
| **Primers for the RNAi constructs** | |
| HvERdj3B_RNAi_F1 | CACCGAGGAGGGGCTGAAGCAGT |
| HvERdj3B_RNAi_R1 no stop | ATACATTCCAGGACCAATTTGA |
| HvERdj3B_RNAi_F2 | CACCAACTGCAGGAATGAAGTTTACCA |
| HvERdj3B_RNAi_R2 no stop | GACCAAATGGTTGTCAAGATG |
| **Primers for EtHAn constructs** | |
| pEDV6-F | TGGGTGGAGGTAAACGAGTC |
| pEDV6-R | AATTTCCATTCGCCATTCAG |
